# PhylteR: efficient identification of outlier sequences in phylogenomic datasets

**DOI:** 10.1101/2023.02.02.526888

**Authors:** Aurore Comte, Théo Tricou, Eric Tannier, Julien Joseph, Aurélie Siberchicot, Simon Penel, Rémi Allio, Frédéric Delsuc, Stéphane Dray, Damien M. de Vienne

## Abstract

In phylogenomics, incongruences between gene trees, resulting from both artifactual and biological reasons, can decrease the signal-to-noise ratio and complicate species tree inference. The amount of data handled today in classical phylogenomic analyses precludes manual error detection and removal. However, a simple and efficient way to automate the identification of outliers from a collection of gene trees is still missing.

Here, we present PhylteR, a method that allows a rapid and accurate detection of outlier sequences in phylogenomic datasets, i.e. species from individual gene trees that do not follow the general trend. PhylteR relies on DISTATIS, an extension of multidimensional scaling to 3 dimensions to compare multiple distance matrices at once. In PhylteR, these distance matrices extracted from individual gene phylogenies represent evolutionary distances between species according to each gene.

On simulated datasets, we show that PhylteR identifies outliers with more sensitivity and precision than a comparable existing method. We also show that PhylteR is not sensitive to ILS-induced incongruences, which is a desirable feature. On a biological dataset of 14,463 genes for 53 species previously assembled for Carnivora phylogenomics, we show (i) that PhylteR identifies as outliers sequences that can be considered as such by other means, and (ii) that the removal of these sequences improves the concordance between the gene trees and the species tree. Thanks to the generation of numerous graphical outputs, PhylteR also allows for the rapid and easy visual characterisation of the dataset at hand, thus aiding in the precise identification of errors. PhylteR is distributed as an R package on CRAN and as containerized versions (docker and singularity).

## Introduction

Supermatrix, supertree and coalescent-based approaches are commonly used in phylogenomics to obtain a species tree from a collection of genes. These methods are similar in their first steps: for a list of taxa of interest, a large collection of single-copy orthologous gene sequences is retrieved and a multiple sequence alignment (MSA) is computed for each cluster of orthologous genes (see von Haeseler 2012 for a comparison of these approaches). The methods then differ by the strategy employed. In the supermatrix approach, MSAs are concatenated into a supermatrix that is used to build a phylogeny, generally with Maximum Likelihood (ML) or Bayesian methods (such as IQ-TREE, Minh, Schmidt, et al. 2020; or Phylobayes, Lartillot et al. 2013). In the supertree and coalescent-based approaches, individual gene trees are built from individual MSAs and a species tree is obtained by combining them all, *e.g.* with MRP (Baum 1992; Ragan 1992; Ronquist 1996), MP-EST (Liu et al. 2010) or ASTRAL (Zhang et al. 2018) to only cite a few.

Regardless of the method employed, errors in the individual gene MSAs and errors in the individual gene trees (leading to incongruences with the species tree) can negatively impact the quality (accuracy) of the reconstructed species tree (Philippe et al. 2017).

For MSAs, various filtering methods have been developed, categorised into two groups: methods that entirely remove sites (columns) or sequences (rows) from the alignment (trimAl, Capella-Gutiérrez et al. 2009; BMGE, Criscuolo and Gribaldo 2010), and methods that are more *picky* and allow identifying and filtering (or masking) small segments in the alignments (Divvier, Ali et al. 2019; HmmCleaner Di Franco et al. 2019; TAPER, Zhang et al. 2021). The latter group of methods was shown to be a better choice for alignment filtering, leading to better gene tree topologies (closer to the species tree) and more consistent terminal branch lengths in gene trees (Ranwez and Chantret 2020).

For collection of gene trees, filtering methods also exist, and the categorization into two groups of strategies still holds. In the first group are methods that prune rogue taxa, *i.e.* taxa that are unstable among gene trees (RogueNaRok, Aberer et al. 2013) and methods that eliminate orthologous gene families whose history is uncorrelated with the others. In the second group are more *picky* methods that identify and filter out only some species in some genes trees (*i.e.* Phylo-MCOA, de Vienne et al. 2012; or TreeShrink, Mai and Mirarab 2018). Just like for the alignment filtering methods seen above, *picky* approaches are thought to provide the best compromise between removing sequences with conflicting phylogenetic signals and keeping the maximum information content.

Filtering (sometimes called trimming) MSAs is now done routinely in phylogenomic pipelines, with methods that can be applied automatically to large datasets (see above). But just because we apply a filter at the MSA level doesn’t mean we shouldn’t filter gene trees also. Some of the reasons that lead to incongruences between gene trees and the species tree (and among gene trees), i.e. gene tree-building errors, undetected paralogy, horizontal gene transfer (HGT) and Incomplete Lineage Sorting (ILS), may not be detectable at the MSA stage. For filtering individual gene trees, reference methods do not exist yet (Philippe et al. 2017). Indeed, identifying species in gene trees whose position is not concordant with their position in the other gene trees (referred to as outliers in de Vienne et al. 2012 and hereafter) is still commonly done by eye (when it is done), which is highly questionable in terms of efficacy, objectivity, and reproducibility.

Here we present PhylteR, a new phylogenomics filtering method that can accurately and rapidly identify outliers in a collection of gene trees. Unlike Phylo-MCOA (de Vienne et al. 2012), from which it is largely inspired, it is an iterative process where obvious outliers are removed first, leaving space for better identification of more subtle ones, and leading *in fine* to a finer identification of outliers. Unlike TreeShrink (Mai and Mirarab 2018), it is not based solely on the diameter of unrooted gene trees and is thus more accurate when outliers are not associated with long branches (e.g. topological incongruences). Also, PhylteR relies on the multivariate analysis method DISTATIS (Abdi et al. 2005; Abdi et al. 2012), which is specifically designed to compare distance matrices, unlike Multiple Co-inertia Analysis (MCOA, Chessel and Hanafi 1996) used in Phylo-MCOA (de Vienne et al. 2012), and is thus more appropriate for the problem at hand We tested PhylteR on two types of datasets: simulated datasets where outliers were known, and a biological dataset comprising 14,463 genes for up to 53 species previously used for Carnivora phylogenomics (Allio et al. 2021). For the simulated datasets, horizontal gene transfers (HGTs) were simulated and recorded, and sequences affected by these HGTs were considered as outliers. The simulations also included various degrees of Incomplete Lineage Sorting (ILS), a phenomenon where within-species polymorphism lasts longer than the time between two successive speciations (Scornavacca and Galtier 2017), leading to incongruences between gene and species trees and among gene trees. The importance of this phenomenon as a source of incongruence in phylogenomic datasets is not clear, but because recent coalescence-based methods are able to handle ILS-induced signal explicitly (e.g. ASTRAL, Zhang et al. 2018), it seemed pertinent to evaluate whether PhylteR was (or not) sensitive to it. In the empirical dataset, outliers were of course unknown but “properties” associated to gene sequences could be gathered (see Shen et al. 2016 for a list of such properties), so that enrichment of sequences having some of these properties in the list of outliers could be quantified. Finally, we looked at the effect of PhylteR on the overall concordance between the gene trees and the species tree after filtering. We compared the results with those obtained with TreeShrink (Mai and Mirarab 2018), the only other tool to our knowledge with a similar objective that could reasonably be applied on such a large dataset.

We show that PhylteR correctly identifies species in gene trees whose phylogenetic placement is not in accordance with its placement in other gene trees, and that this holds even in the presence of incongruence among gene trees due to ILS. We also provide strong evidence that the automatic removal of outliers with PhylteR improves the concordance between gene trees and the species tree in greater proportions than TreeShrink (Mai and Mirarab 2018).

We hope that PhylteR could become the standard that was lacking (Philippe et al. 2017) for cleaning datasets prior to species tree reconstruction in phylogenomic pipelines.

## Material and Methods

### Description of the PhylteR method

The PhylteR method, in its entirety, is depicted in Figure 1. It starts with *K* distance matrices obtained from *K* genes by computing pairwise distances (sum of branch lengths) between species in each gene tree. All the matrices are given the same dimensionality by filling missing data (if any) with the mean value across matrices, and are then normalised by dividing each matrix by either its median or its mean value (default is median). The normalisation by median prevents genes from fast- and slow-evolving orthologous genes to be erroneously considered as outliers, and appears as a better choice than a normalisation by the mean as it is less affected by outlier values.

**Figure 1.**
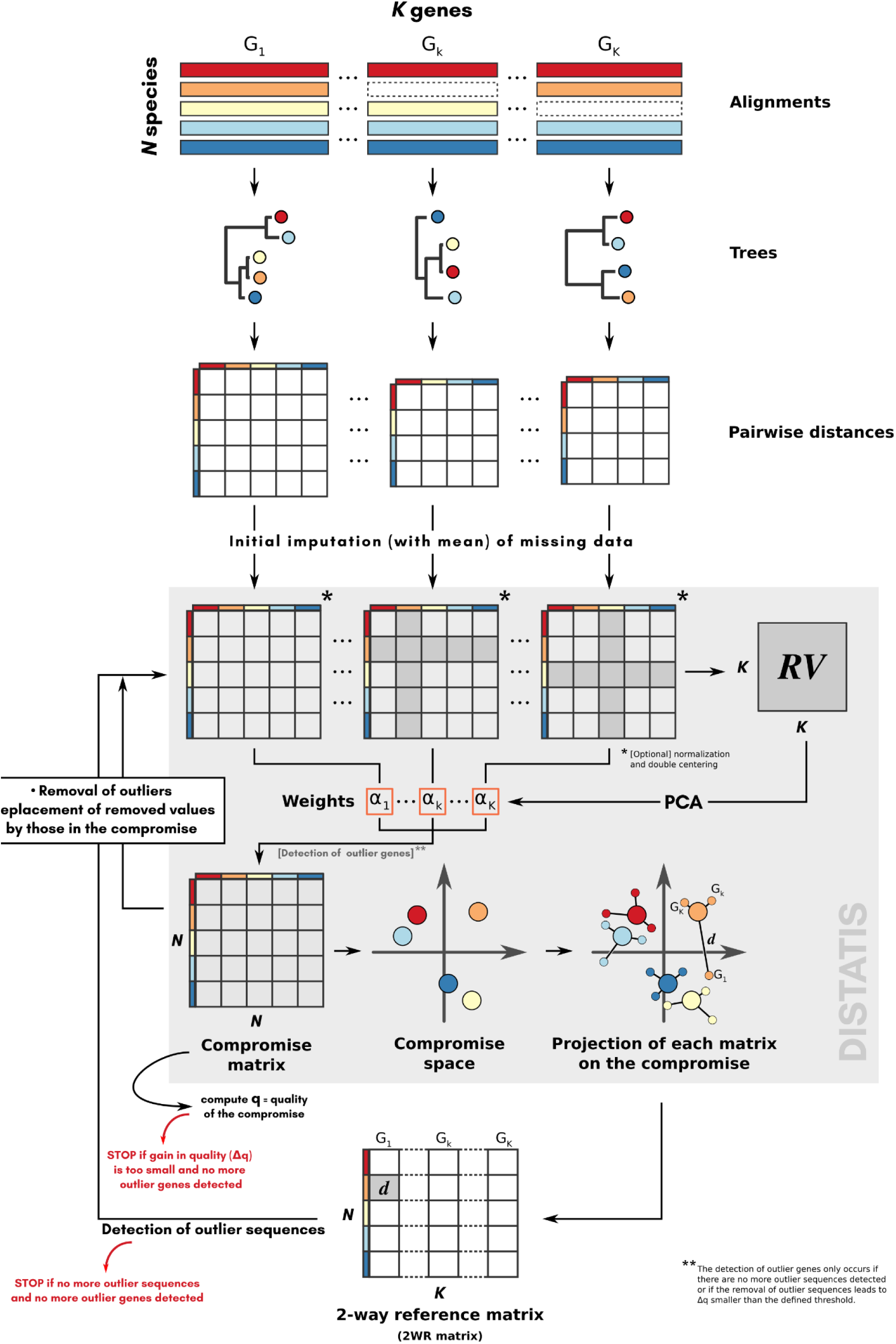
Principle of the PhylteR method for identifying outliers in phylogenomic datasets. The method relies on DISTATIS (grey block), an extension of multidimensional scaling to three dimensions. See text for the detail of the different steps.

From the *K* matrices obtained, an incremental process starts consisting in three main steps detailed in the next sections: (1) comparison of the matrices with the DISTATIS method (Abdi et al. 2005; Abdi et al. 2012), (2) detection of outliers sequences, and (3) evaluation of the impact of removing these outliers on the overall concordance between the matrices. Note that we refer to *outlier sequence* as a single gene for a single species (one sequence in one alignment, or one tip in one gene tree) that does not follow the general trend (i.e. other alignments or gene trees), while *outlier gene* refers to a complete alignment (or a complete gene tree) that does not agree with the other alignments (or gene trees).

These steps are repeated until no more outlier sequences are detected, or until the removal of the identified outlier sequences does not increase the concordance between the matrices more than a certain amount specified by the user. Before finishing the optimization, PhylteR performs a last action consisting of checking whether some outlier genes still exist despite the removal of outlier sequences already performed. These outlier genes correspond to single-copy orthologous genes for which the lack of correlation with others is not due to a few outlier sequences but are globally not following the trend. If outlier genes are discarded there, the optimization restarts as it may have unblocked the detection of other outliers.

### Comparison of individual gene matrices with DISTATIS

DISTATIS is a multivariate method designed to evaluate the concordance between *K* distance matrices (*K* orthologous genes) measured on the same *N* species. The principle of DISTATIS is depicted in Figure 1 (grey box). The first step of DISTATIS consists of computing a matrix of RV coefficients (Robert and Escoufier 1976) that measures the similarities between the species pairwise distances present in each matrix. This can be seen as an extension of the correlation matrix (used in principal component analysis) that, instead of measuring the links between a set of variables, evaluates the relationships between a set of tables (gene distance matrices here). In a second step, a compromise distance matrix is built as the average of the *K* distance matrices weighted by the first eigenvector of the matrix of RV coefficients. The compromise represents the best consensus between the *K* distance matrices, as the weights used in the averaging procedure take into account the similarities between them (i.e., more similar distance matrices would have more weights in the definition of the compromise). In a third step, the compromise matrix is submitted to an eigendecomposition procedure so that species can be represented in a low-dimensional multivariate space. In this compromise space, species are positioned so that their distances (computed in few dimensions, see after) represent the best approximations of the original distances contained in the compromise matrix. We used a broken stick model (Barton and David 1956) to estimate the number of dimensions (axes) of the compromise space, as this simple method was shown to give a good approximation of the correct dimensionality of the data with another multivariate approach (Jackson 1993). Then, each individual pairwise distance matrix is projected on the compromise space. This allows us to obtain a representation of species associated with each gene family. In other words, the compromise identifies the dissimilarities between species that are common for all genes whereas the projections of individual distance matrices allow depiction of the peculiarities of each sequence. Lastly, we compute the distances, in the compromise space, between the position of a species given by all genes (the compromise) and its position associated to a particular gene family (using the projection procedure) and filled a gene x species 2-Way Reference matrix (2WR matrix, see figure 1) with these values.

### Detection of outlier sequences from DISTATIS results

From the 2-Way Reference matrix (2WR matrix, see figure 1), we apply the method of Hubert and Vandervieren (2008) to detect all values that are outliers, at the right of the univariate distribution of values. This method is an adjustment of the Tukey method (the classical boxplot) adapted to skewed distribution. In brief all values above

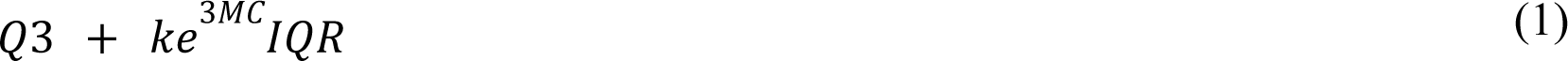

are considered outliers. *Q3* is the 3rd quartile of the distribution, *IQR* is its interquartile range and *MC* is the medcouple of the distribution (Brys et al. 2004), a measure of skewness bounded between -1 (left skewed) and +1 (right skewed). The *k* value is chosen by the user (default is 3), and controls how stringent the detection of gene outliers is. Small values of *k* lead to more gene outliers being detected. The detection of gene outliers is performed after normalisation of the 2WR matrix, achieved by dividing each row (the default) or each column by its median. This normalisation leads to an exaggeration of outlier values, easing their identification.

### Detection of outlier genes

When no more outlier sequences are found in the 2WR matrix, PhylteR checks whether some genes are still uncorrelated to others. These outlier genes are detected by finding outlier values in the weight array (*α_1_, α_2_, …, α_K_*, see Figure 1). The outlier detection method used is the same as for the outlier sequences of the 2WR matrix (Equation 1) but its stringency can be tuned independently (with parameter *k2* in place of parameter *k* in Equation 1, defaulting to *k2* = *k* = 3).

### Exit criteria of the PhylteR iterative process

PhylteR is an iterative process (see Figure 1) with two exit points. The first one is straightforward: if no more outlier sequences are detected in the 2WR matrix, and if no more outlier genes exist (see above), then the process stops. The second one is based on the gain (Δ*_q_*) achieved by removing outlier sequences (i.e. the change in *q*, the quality of the compromise). If this gain is below a certain threshold (10^-5^ by default), and if no more outlier genes exist, then the process stops.

### Evaluation of the PhylteR method

#### Datasets

We used three types of datasets to evaluate PhylteR and compare it with TreeShrink: a simple dataset used for illustrative purpose only, a collection of simulated examples obtained with the program SimPhy (Mallo et al. 2016), and a large Carnivora phylogenomic dataset with 53 species (Allio et al. 2021). These datasets are described below.

● *Simulated dataset for illustrative purpose:* we generated a small collection of gene trees in order to illustrate the different steps of the PhylteR process. A single phylogenetic tree with 20 species was randomly generated with function `rtree()` from package *ape* v5.6.2 (Paradis and Schliep 2019). This tree was duplicated 25 times to mimic 25 orthologous gene families. To add variance to branch lengths, a value sampled in a normal distribution with mean 0 and standard deviation *0.15* was added to each branch length of each tree (if the resulting branch length was negative its absolute value was taken). Ten outliers were then generated by randomly sampling 10 times a species in a gene tree and moving it to another random location.
● *Simulated datasets: We* simulated collections of gene trees with known outliers in order to evaluate PhylteR and compare it with TreeSkrink. We used SimPhy (Mallo et al. 2016), a program that can simulate the evolution of gene families (and thus gene trees) given a species tree under various evolutionary processes including HGT but also ILS. We used, as a species tree, the 53-taxa carnivora tree of Allio et al. (2021), the same as for the biological dataset (next point). To be usable in SimPhy, we transformed the tree to ultrametric with function *chronos* in ape (Paradis and Schliep 2019) and we rescaled the branch lengths so that the root-to-tip distance reflected (roughly) the number of generations, *i.e.* 8,899,579 generations in this case. This value was obtained by dividing the age of the root of the tree (74 millions years old, (Kumar and Subramanian 2002)) with a rough estimate of the generation time in carnivora (8.315 years if taking the median of the generation times of the species studied in (Kerk et al. 2013)).

For each replicate (100 each time), collections of 500 gene trees were simulated by setting the rate of HGT to 1e-8, the tree-wide substitution rate to 2.2e-9, and varying the level of ILS by changing the population size: 10 (NO-ILS), 100,000 (LOW-ILS), 200,000 (MODERATE-ILS), 500,000 (HIGH-ILS; the detailed commands used for Simphy are given as supplementary method). Then, from the 500 trees obtained, only 100 were retained, randomly sampled among those where at most one horizontal gene transfer occurred (to allow unequivocal identification of outliers), and for which the transfer (if any) changed the topology of the gene tree. The whole process was repeated three times, varying the maximum number of outliers allowed per gene, between 1, 10 and 53 (theoretical max). This allowed exploring the impact of the number of outliers per gene (linked here to the age of the HGT) on the capacity to correctly identify outliers, *i.e.* the species that were changing position relative to the species tree because of HGT. For ILS, it was not possible to identify precisely what species should be considered as outliers or not. We could only look at the impact of ILS on the mean topological distance between the collection of gene trees and the species tree (mean RF distance between 3 for NO-ILS and more than 20 for HIGH-ILS, see Figure S1) and evaluate whether this had an impact or not on the precision and sensibility of our method.

- ● *Carnivora dataset (CD):* We used the raw sequence files (before alignment and filtering) from a previously assembled phylogenomic dataset comprising 14,463 genes for 53 species aimed at resolving the phylogeny of the order Carnivora (Allio et al. 2021). This dataset was obtained by extracting single-copy protein-coding orthologous genes from the genomes of 52 carnivore species, plus the Malayan pangolin (*Manis javanica*) used as outgroup, following the orthology delineation strategy of the OrthoMaM database (Scornavacca et al. 2019). These raw sequence files were aligned and filtered using the OMM_MACSE pipeline (Ranwez et al. 2021), which combines (i) translated nucleotide sequence alignment at the amino acid level with MAFFT (Katoh and Standley 2013), (ii) nucleotide alignment refinement (based on amino acid alignment) with MACSE v2 (Ranwez et al. 2018) to handle frameshifts and non-homologous sequences (Ranwez et al. 2018), and (iii) masking of ambiguously aligned and dubious parts of sequences with HMMcleaner (Di Franco et al. 2019). In the original study (Allio et al. 2021), this Carnivora dataset was successfully filtered using an early version of PhylteR allowing the removal of outlier sequences and genes generating abnormally long branches. Therefore, it was a good candidate dataset to test the completely redesigned and improved version of PhylteR presented here.

#### Evaluation of the accuracy of PhylteR outlier detection and comparison with TreeShrink

We evaluated PhylteR’s ability to detect outliers that are either correct (when it is possible to test it, with simulated datasets) or meaningful according to the biological information we can gather from the dataset at hand.

We used the first simulated dataset for illustration purposes only. For the other simulated datasets, *i.e.* for each level of ILS, for different maximum numbers of outlier species per gene (1, 10 and 53) and for each one of the 100 replicates, we ran PhylteR with default parameters and we counted the number of True Positives (TP, outliers that were simulated and that are retrieved), False Positives (FP, outliers that were not simulated but are identified) and False Negative (FN, outliers that were simulated but are not retrieved). From those, we computed the mean precision (TP/(TP+FP)) and recall (or sensitivity, TP/(TP+FN)) of the outlier identification of PhylteR. An estimate of the expected precision and recall when the same number of outliers were randomly sampled was also computed. To evaluate the impact of ILS-induced incongruences between gene trees on the ability of PhylteR to correctly identify outliers, precision and recall were computed and compared between the four levels of ILS simulated (NO-ILS, LOW-ILS, MODERATE-ILS and HIGH-ILS). Finally, for comparison purposes we performed the same analyses using TreeShrink v1.3.9 (Mai and Mirarab 2018) in place of PhylteR with default parameters for detecting outliers.

For the Carnivora dataset, we have no access to the *true* outliers. It is thus impossible to compute precision and recall on this empirical dataset as done on the simulated ones. Instead, we can compute “features” associated to each gene sequence for each species (*sequence* hereafter), that are, *a priori*, associated with errors or with lack of signal in phylogenomic datasets. We can then evaluate whether the outliers detected by PhylteR are enriched in extreme values for these features, as compared with randomly selected sequences or with outliers identified with TreeShrink. The list of features and the reason for their choice is listed below.

● **Sequence length:** Long sequences were shown to carry more phylogenetic signal than shorter ones (Salichos and Rokas 2013; Shen et al. 2016). To explore the possible enrichment of outliers in short sequences, we computed the length (in bp) of each sequence in each gene MSA, and explored its distribution in outliers.
● **Duplication score**: when a sequence in a gene tree is not orthologous to the others but is a paralog, its localization in the gene tree is likely to be incorrect. To have an insight into the level of “paralogousness” of each sequence in the Carnivora dataset, we compared the Carnivora species tree published in Allio et al. (2021) with each one of the 14,463 gene trees using the reconciliation program *ALEml_undated* (Szöllősi et al. 2015). This tool allows inferring the duplications, losses and transfers experienced by a gene by comparing its history (the gene tree) with that of the species (the species tree). Here we inferred only duplications and losses (transfer rate was forced to be 0), we forced the origination of each gene at the root of the species tree (parameter O_R=10000) and we used default values for all other parameters. We then computed the number of duplications inferred from the root to each tip of each gene tree, and normalised this value by the number of nodes encountered. This value represents the normalised number of duplications experienced by each sequence, whose distribution in outliers could be evaluated.
● **Hidden paralogy, the KRAB Zinc finger (KZNF) protein family case:** The KZNF super-family is actively duplicating in vertebrates with hundreds of paralogs per genome (Huntley et al. 2006; Liu et al. 2014). Thus, the orthologous relationships between these proteins is expected to be hard to retrieve and the reconstructed orthologous gene families are likely to contain hidden-paralogs. If an outlier detection method is indeed able to remove hidden paralogs, we should see an enrichment of KZNF genes in the list of outliers.
● **Synteny:** Synteny (in our sense) is the link between two genes occurring consecutively on a genome, *i.e.* without any other gene (in the dataset) located between them. One gene then has two synteny linkages. A synteny *break* occurs when two genes are consecutive in one species but their orthologs in another species are not. The direction of transcription (coding strand) is considered, *i.e.* if it has changed it is considered as a break even if the genes appear in the same order. One gene, compared to its ortholog in another species, may then be associated with 0, 1 or 2 breaks. We call genes associated with 2 breaks *syntenic outliers*. We test if outliers found by PhylteR are more often syntenic outliers than randomly sampled genes. Our rationale behind this question is that synteny breaks are due to genomic rearrangements (inversions, duplications, translocations, …), but can occur in the data, and in much larger proportion, for many artifactual reasons: annotation errors, assembly errors, or orthology assessment errors. These different sources of errors are expected to lead to phylogenetic placement errors for the species carrying the affected genes. We thus formulate the hypothesis that outlier genes may be more often associated with synteny breaks than randomly sampled genes. To evaluate this, we focused on 14 Carnivora genomes (Table S1) that we compared in a pairwise manner. For each pair we compared the list of syntenic outliers with the list of outliers retrieved by each outlier method tested, and we computed the p-value associated with the observed size of the intersection under the hypothesis that the two sets of outliers are independent.

In order to compare the distributions of values for the different features listed above between outlier detection methods, we needed lists of outliers of comparable size. The number of outliers retrieved with default parameters being very different with the two methods using default parameters (7,183 with PhylteR *vs* 19,643 with TreeShrink, see Table 2), we created two collections of outliers, a **small** and a **large** one (Table 2). For the **small** collection, we selected a value for the parameter *q* in TreeShrink in order to get a number of outliers as close as possible to the number of outliers obtained with PhylteR default parameters. This was achieved for *q* = 0.012, leading to 7,032 outliers. For the **large** collection, we selected a value of the *k* (and *k* = *k2*) parameter in PhylteR leading to a number of outliers as close as possible to the number of outliers detected with TreeShrink default parameters. This was achieved for *k* = 1.55, leading to 20,157 outliers. Parameters used and number of outliers in each collection and with each outlier detection method are presented in Table 2.

**Table 2.** Collections of outliers used to evaluate PhylteR and compare it to TreeShrink. The **small** collection is obtained by tuning the TreeShrink parameters in order to obtain roughly the same number of outliers as with the default parameters of PhylteR. The **large** collection is obtained in the opposite way.

| Collections | PhylteR |  | TreeShrink |  | Random |
| --- | --- | --- | --- | --- | --- |
|  | Parameters | # outliers | Parameters | # outliers | # outliers |
| **small** | *default* | 7,183 | $q = 0.012$ | 7,032 | 7,183 |
| **large** | $k = k2 = 1.55$ | 20,157 | *default* | 19,643 | 20,157 |

#### Evaluation of the impact of outlier sequences removal on species tree support

It is expected that a tool that accurately removes outliers in phylogenomic datasets should increase the concordance between the gene trees and the species tree. To evaluate this and compare PhylteR with randomly sampled sequences and with TreeShrink-identified outliers, we computed the gene concordance factor (gCF, Minh, Hahn, et al. 2020) as implemented in IQ-TREE version 2.1.3 (Minh, Schmidt, et al. 2020) for every branch in the Carnivora species tree (obtained from Allio et al. 2021). For each branch of the species tree, this factor indicates the percentage of gene trees in which this branch is found (among gene trees where this can be computed, or “decisive” trees, see Minh, Hahn, et al. 2020). gCF was computed according to either the original gene trees (gCF_init_), or to a list of gene trees obtained after pruning outliers (four sets of gene trees corresponding to the four list of outliers in Table 2).

In order to see the effect of outliers removal on the concordance factor, we computed the difference (ΔgCF) between gCF_init_ and every other gCF, separating the small and the large collections of outliers. Positive values of ΔgCF indicate that a branch is more supported after filtering than before. Comparing ΔgCF between PhylteR and TreeShrink gives an indication of whether, for the same total number of outliers removed, PhylteR performs better than TreeShrink at identifying sequences with conflicting phylogenetic signals and increasing the concordance between the species tree and the gene trees.

## Results

### Illustration of the general principle of PhylteR

The different steps of the PhylteR process (Figure 1) are illustrated on a simple example dataset comprising 25 genes for 20 species, with 10 outliers. The main steps are as follows. Individual gene trees are transformed into individual gene matrices that are then combined into a unique *compromise* matrix obtained after weighting each matrix by its concordance with the others: matrices that are poorly correlated with the others have less weight in the creation of the compromise (Figure S3A-E). This matrix is then projected onto a space on which individual matrices are projected as well (Figure 2A and S3F). By computing the distance of each species in each orthologous gene to its reference position in this projection, the two-way reference matrix is obtained (Figure 2B and S3G). It is from this matrix that outlier sequences can be identified and removed.

**Figure 2.**
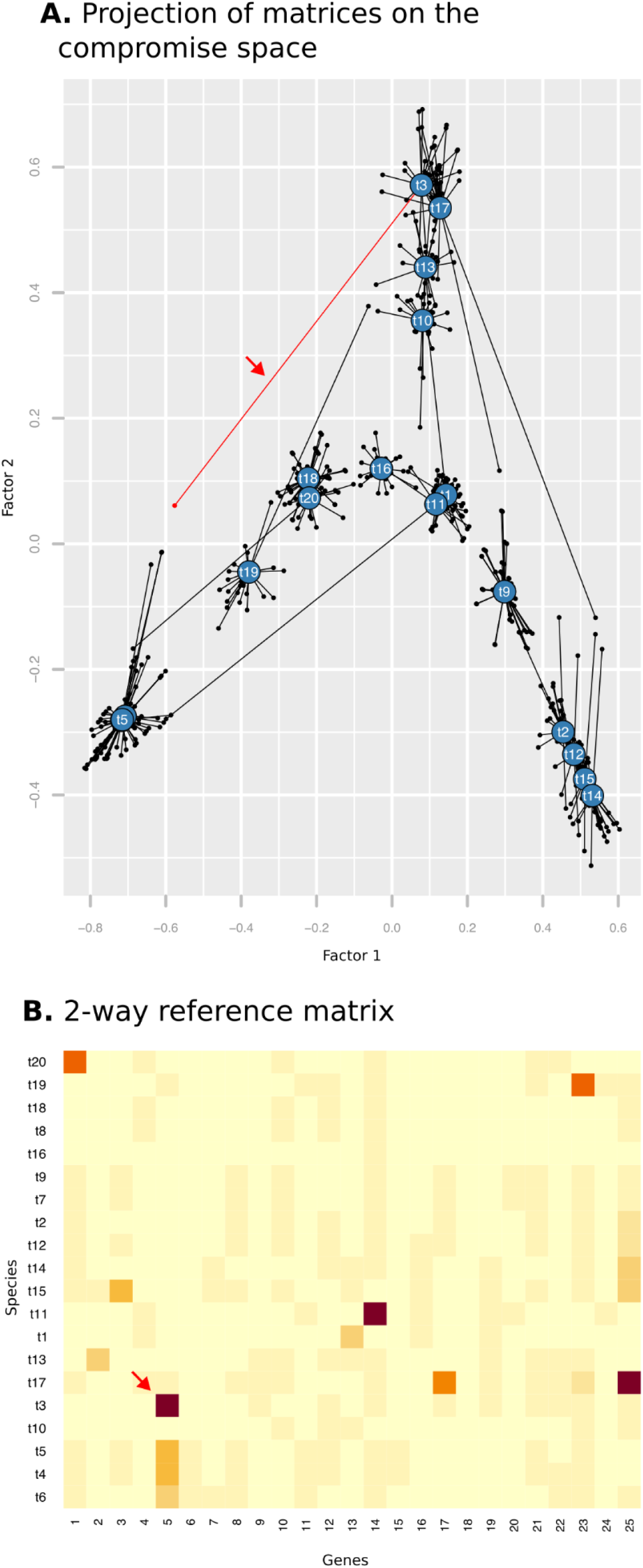
Two objects of the PhylteR process. **A:** the compromise matrix is projected into a multidimensional space (the two first axes only are represented here). This gives the reference position of each species relative to each other (blue badges with species names on it). Individual gene matrices are projected on the same space (small dots) and the distance between each gene in each species to its reference position is represented by a line. The red line and the red arrow identify species t3 in gene 5. This projection is transformed into a 2D matrix (**B)** by computing the distance between each species in each gene to its reference position (i.e. the length of each line in **A**). The gene ✕ species matrix obtained, that we refer to as the 2-way reference matrix (2WR) is used to detect outliers like the one indicated by the red arrow, corresponding to the red arrow in **A**.

### PhylteR performs well on simulated examples and is robust to ILS-induced incongruences

To evaluate the precision and sensitivity of PhylteR, we used it on four simulated datasets with increasing levels of ILS (NO-ILS, LOW-ILS, MODERATE-ILS and HIGH-ILS). We also computed precision and recall on the same datasets using another method, TreeShrink (Mai and Mirarab 2018). Finally, we computed the expected precision and sensitivity if the same number of sequences identified as outliers by PhylteR and TreeShrink were randomly selected from all sequences.

The outliers that we considered were single species whose position was moved to a new location in some gene trees because of HGTs. For this type of outliers, we observe that PhylteR performs well, precision and recall being close to their maximum value 1 (Figure 3). On the other hand, TreeShrink performs badly, identifying few correct outliers (leading to a mean precision close to 0), but still detecting a large collection of false positives (leading to a low sensitivity). When increasing the maximum number of outlier species in each gene tree to 10 (Figure S2A) or to 53 (Figure S2B), we observe that both the precision and sensitivity of PhylteR slightly decrease while the ones of TreeShrink increase, but the difference between both remains in clear advantage of PhylteR (in this specific setting).

**Figure 3.**
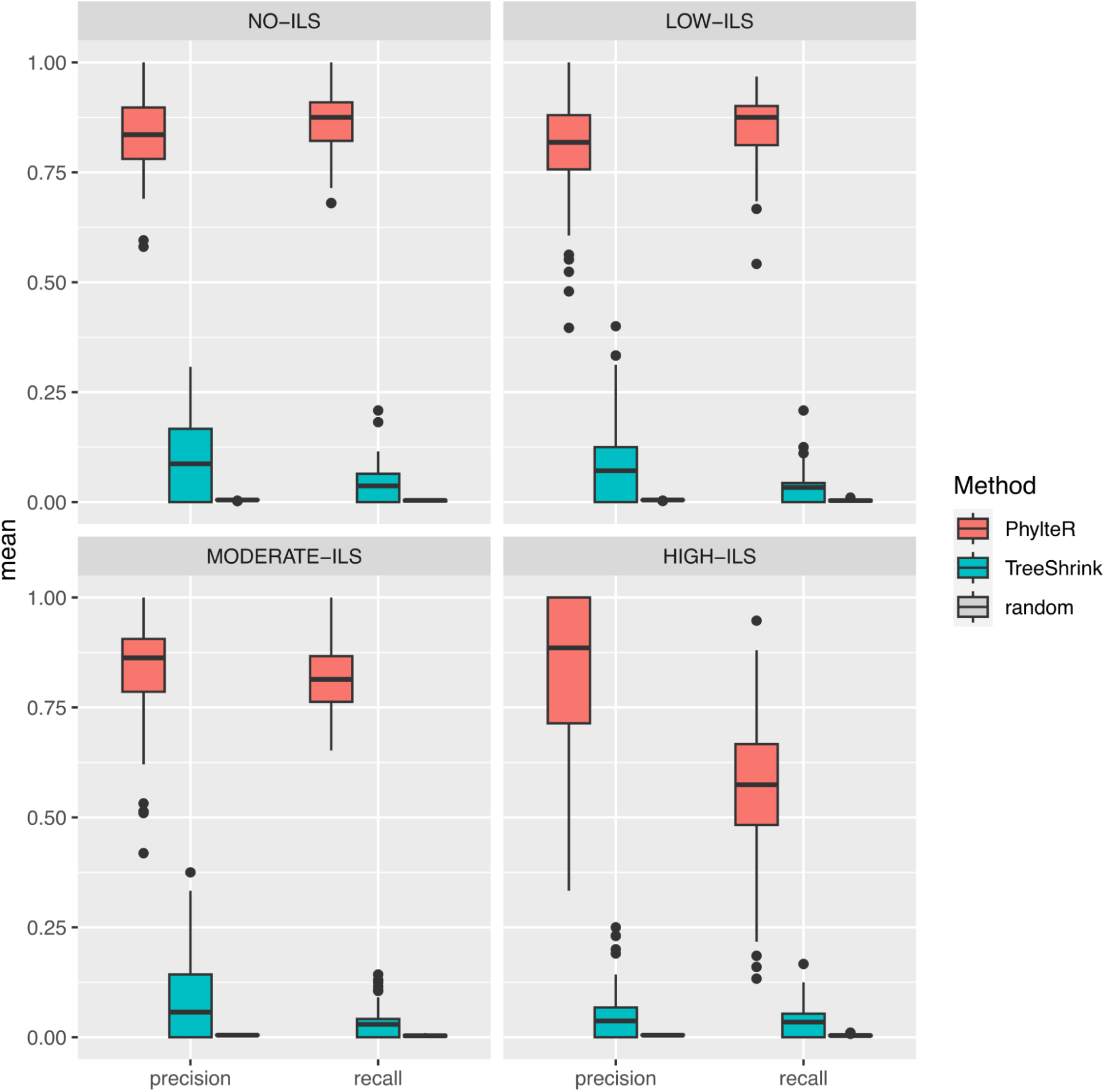
Comparison of the precision and recall (or sensitivity) of the PhylteR and the TreeShrink outlier detection methods for four conditions of Incomplete Lineage Sorting (ILS).

Of note, the level of ILS has almost no effect on the precision and sensitivity of the PhylteR (and TreeShrink, even though negative effect would be hard to see when starting from such low precision and sensitivity values), except when reaching very high ILS (bottom-right panels in Figure 3 and Figure S2A and S2B). In other words, even when the mean topological distance between the gene trees and the species tree is multiplied by more than 5, as is the case between the NO-ILS and the MODERATE-ILS conditions (Figure S1), the precision and sensitivity of PhylteR for detecting the outliers simulated by HGTs do not decrease. This suggests that PhylteR does not consider species that have changed position in some gene trees due to ILS as outliers. This apparent robustness of PhylteR to ILS can be seen as a desirable feature, e.g. when using species tree reconstruction tools that explicitly handle ILS such as ASTRAL (Zhang et al. 2018).

### Characterisation of outliers detected with PhylteR on the Carnivora dataset

Outliers in phylogenomic datasets can be of different nature: fast or slow evolving genes in some species, leading to respectively long or short branches in gene trees, or species being placed in aberrant position in some genes because of horizontal gene transfers (HGT), hidden paralogy, saturated signal, compositional bias, long-branch attraction, or other artifactual reasons (Schrempf and Szöllősi 2020).

In the set of 14,463 gene trees analysed by PhylteR, two sets of outliers (7,183 and 20,157 sequences) were identified with PhylteR (with default or tuned parameters, respectively) and 7,032 and 19,643 with TreeShrink (with tuned and default parameters respectively, see Table 2). A simple comparison of the list of outliers of similar sizes revealed that the overlap between the two lists of outliers was quite small (around 20%, Figure 4). This corresponds to about 70% of the outliers detected by PhylteR being absent from the list of outliers detected by TreeShrink, and vice versa. This reveals fundamental differences between the two approaches.

**Figure 4.**
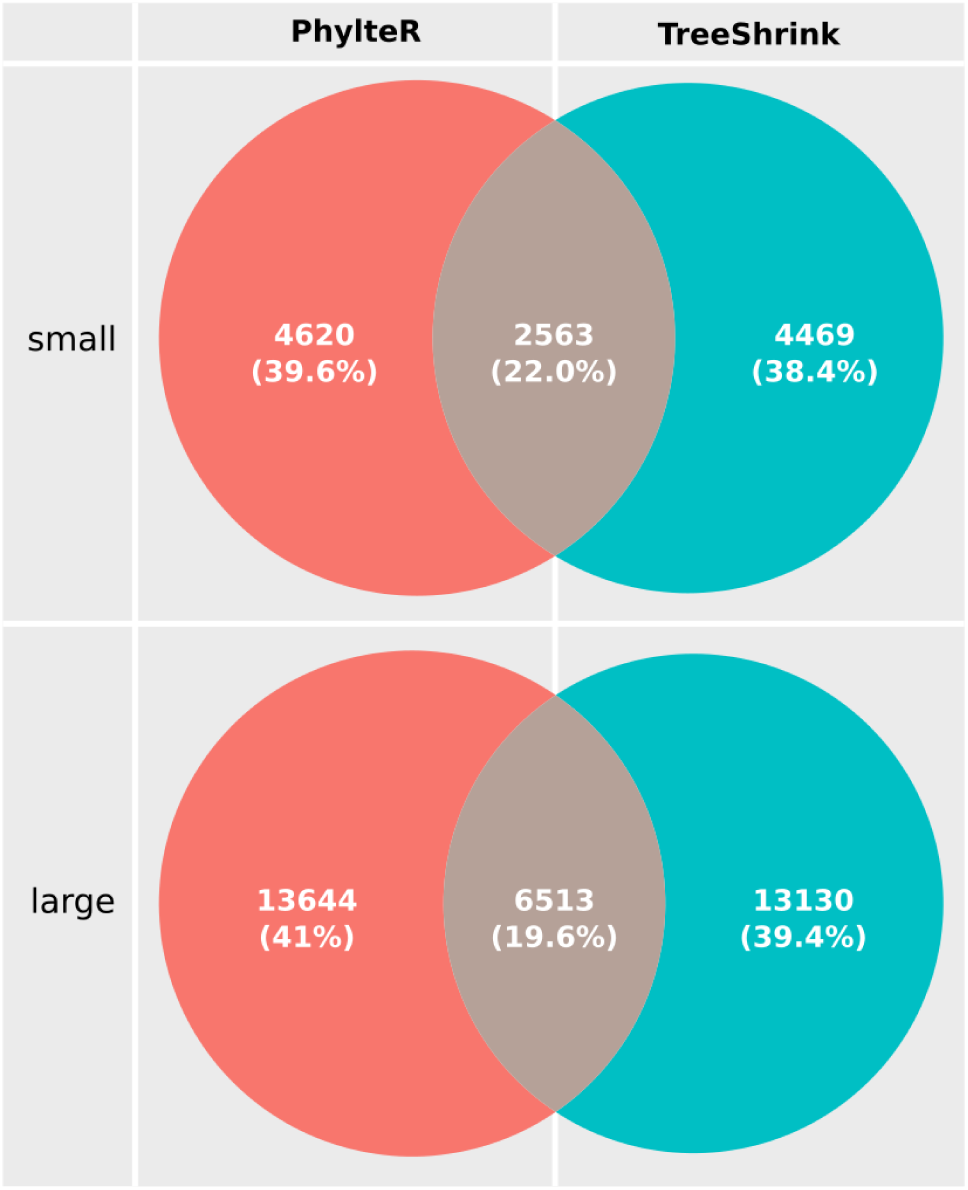
Comparison of the sets of outliers detected by PhylteR (left column) and TreeShrink (write column) on the Carnivora dataset. The two collections of outliers (small and large) correspond to different stringency for the detection of outliers (see Table 2).

To better understand what differs between the outliers detected by PhylteR and those detected by TreeShrink, we compared the distribution values of different features describing these outlier sequences.

First, we observed a significant decrease in sequence length in outlier sequences for both PhylteR and TreeShrink as compared to randomly sampled sequences (p<2.2e-16 in both cases and for both collections of outliers, Figure 5A). Sequence lengths were higher in PhylteR outliers than in TreeShrink outliers for the small collection of outliers (p<2.2e-16) but the opposite was observed for the large collection of outliers (p<3.17e-14). The fact that outliers are enriched in short sequences is thought to be due to the expected correlation between the size of a sequence and the phylogenetic signal it carries. Shorter sequences are more prone to misplacement in phylogenetic trees.

**Figure 5.**
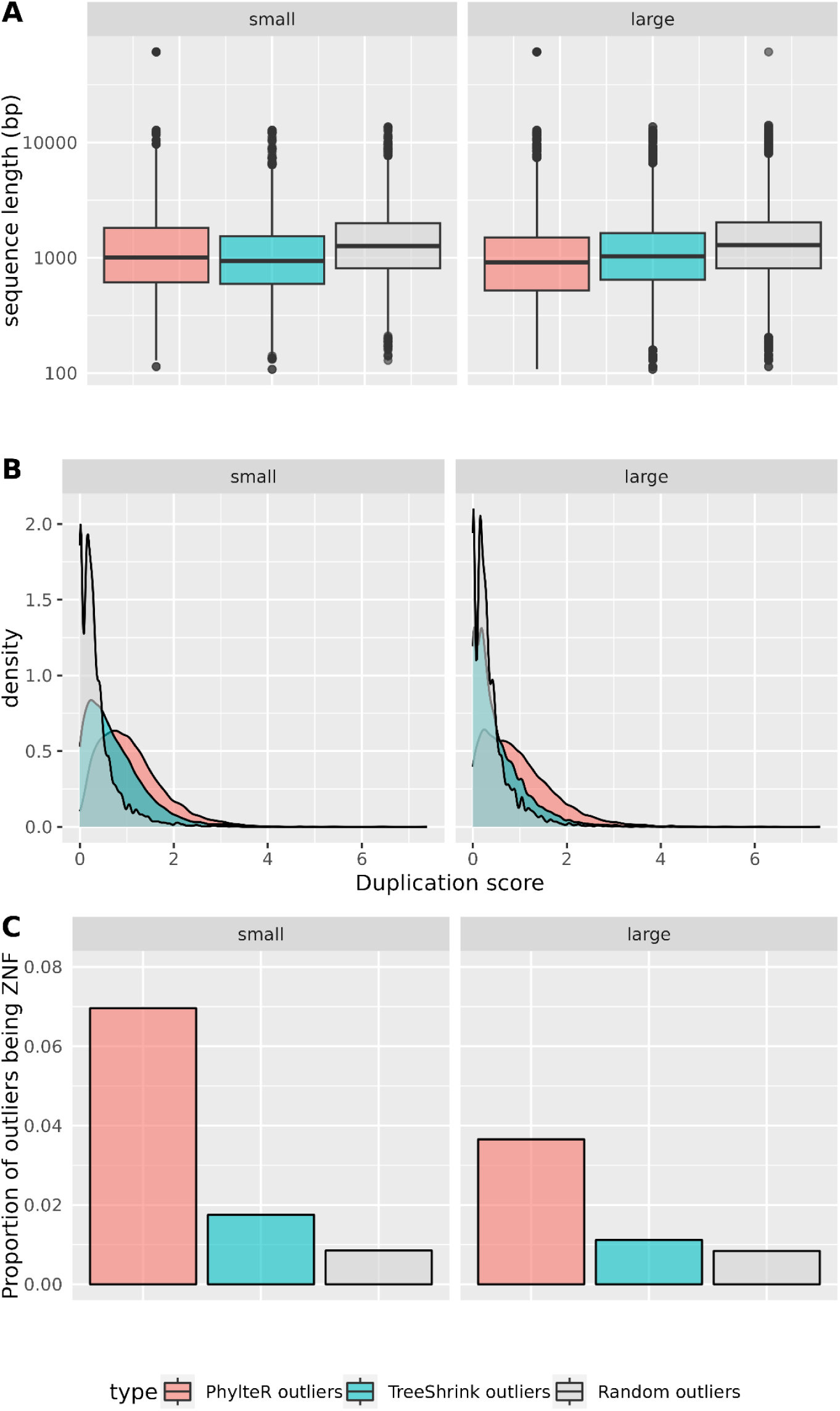
Comparison of distribution values between outliers detected by PhylteR, by TreeShrink, or randomly sampled, for three features associated with outlierness in phylogenomic datasets. A. Distribution of the length (in bp) of the sequence outliers identified by each method. A log scale is used for the y-axis. **B.** Distribution of duplication scores (normalised number of duplications experienced by each sequence) for the outliers identified by each method. **C.** Proportion of outliers being members of the KRAB-ZNF protein family for the outliers identified by each method. The two collections of outliers (small and large) are compared in each case (left and right on each panel).

Second, we compared the distribution of duplication scores in the list of outliers produced by PhylteR and TreeShrink (Figure 5B). We observed a clear difference, for both the small and the large collections of outliers between PhylteR outliers and random outliers, but also between PhylteR outliers and TreeShrink outliers: outliers identified by PhylteR are significantly enriched in sequences that display a higher number of duplications as compared to random or TreeShrink outliers (p<2.2e-16 for all comparisons).

This result is in accordance with the results obtained on simulated datasets: PhylteR is good (and much better than TreeShrink) at identifying misplaced species in some gene trees, which is indirectly what the duplication score captures.

One illustration of the difference between PhylteR and TreeShrink in their ability to capture duplicated sequences (and thus probably hidden paralogues) can be given by the study of peculiar proteins, such as the Zinc-finger family (ZNF). This large family of paralogs first duplicated from the gene PRDM9 or PRDM7 in the ancestor of vertebrates (Emerson and Thomas 2009). These genes are involved in the repression of transposable elements and are still actively duplicating. The high number of duplications renders the resolution of the orthology relationship in this gene super-family very challenging. In the Carnivora dataset, the ZNF super-family has been splitted in 168 orthologous gene families (Allio et al. 2021). As expected in case of hidden paralogy, we see an overrepresentation of the genes belonging to these families in the list of outliers, especially in the outliers identified by PhylteR (Figure 5C). Between 3.79% (for the large set) and 7.4% (for the small set) of PhylteR outliers belong to the ZNF family, while these values drop to 1.78% and 1.12% respectively for TreeShrink outliers, and less than 1% for randomly selected sequences (Figure 5C).

Third, we compared two by two 14 Carnivora species and identified syntenic outliers (see material and methods). In almost all pairwise comparisons, we found that these syntenic outliers significantly overlap the outlier sequences detected by PhylteR. For example, in the comparison between *Zalophus californianus* and *Suricata suricatta* (illustrated in Figure 6), out of the 5,123 genes common to both species in the dataset, 131 (2.56%) are syntenic outliers (i.e. surrounded by two breaks). In comparison, out of the 47 outlier sequences identified by PhylteR (small list) in either *Zalophus californianus* or *Suricata suricatta*, 38 are syntenic outliers (80.8%), which is significantly more than expected by chance (p-value = 1.5e-43). With TreeShrink (small list) for the same pair of species, only 18.1% (17 out of 94) outlier sequences are syntenic outliers, which is much less than with PhylteR but is still significantly different from what is expected by chance (p-value = 1.36e-10). Similar results were obtained for most of the other pairs of species compared (Figure S4 and Supplementary Tables S2 and S3).

**Figure 6:**
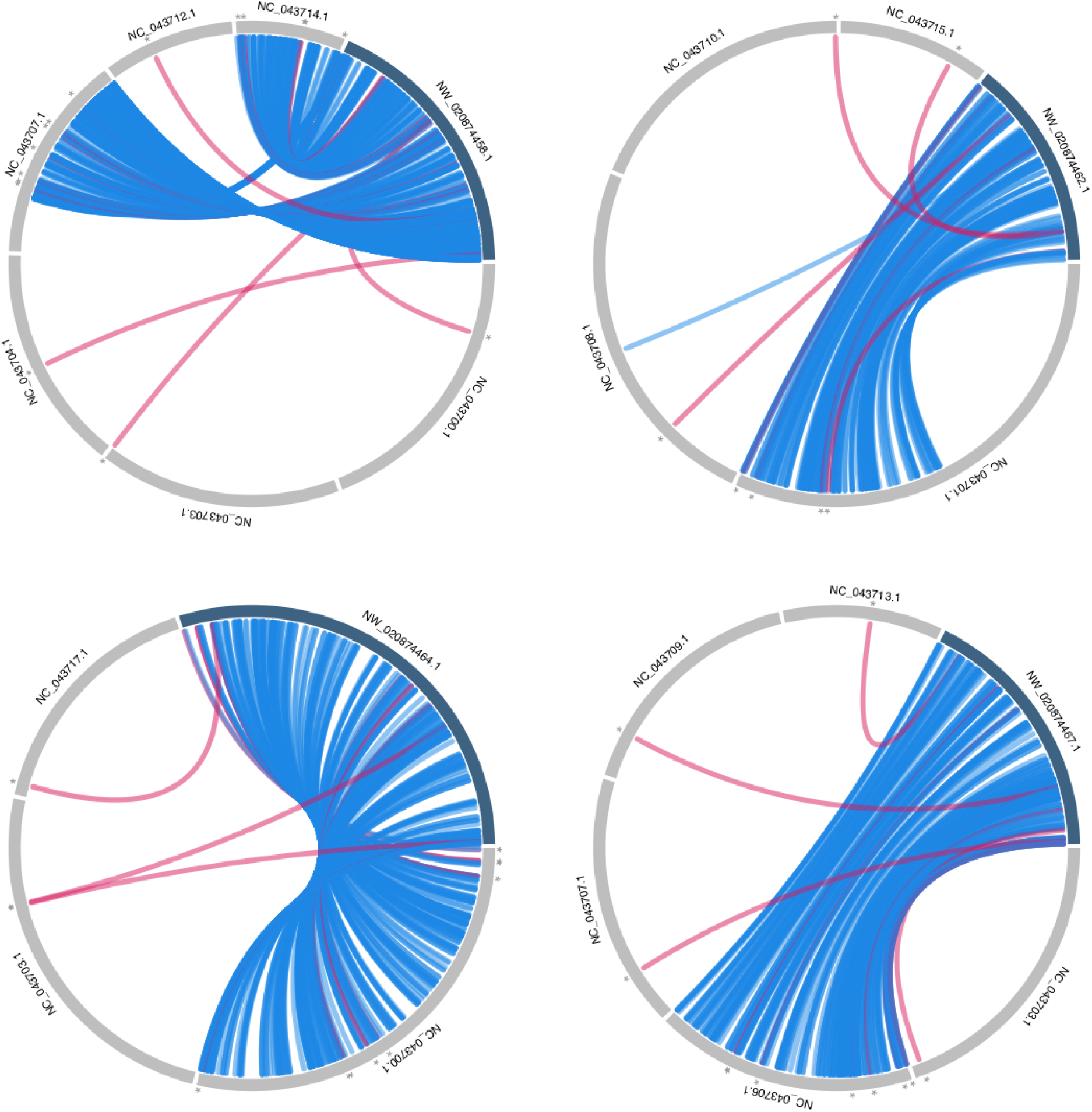
Illustration of the non-syntenic nature of many outliers identified by PhylteR. We represent the comparison of *Zalophus californianus* with *Suricata suricatta* genomes, with *Zalophus* as a reference (arbitrarily, most other pairs of species give similar results). On each circle, a reference *Zalophus* scaffold is represented in dark blue, and all scaffolds for which at least one gene has an ortholog in this scaffold are in light grey. Lines between these scaffolds represent couples of genes annotated as orthologous. Red lines highlight gene outliers detected in *Suricata suricatta*. We observe that they are very often “isolated” genes, *i.e.* syntenic outliers. These genes are thus probably erroneously annotated, erroneously assembled, and their orthology is likely erroneous.

### Impact of filtering outliers on Species Tree support

The gene concordance factor (gCF) is a measure, for a species tree, of how much each one of its branches is supported according to a collection of individual gene trees. A value of 100% means that 100% of the gene trees for which the comparison could be done (“decisive” gene trees in Minh, Hahn, et al. 2020) contain this branch.

Non-random outlier removal processes are expected to increase gCF scores by discarding sequences representing species in gene trees whose position is not in accordance with their placement in the other gene trees. We looked at the difference in gCF score before and after pruning outliers (ΔgCF) for each branch of the Carnivora species tree. For both PhylteR and TreeShrink, an increase in gene concordance was observed. It was higher with PhylteR than with TreeShrink, indicating a better identification of misplaced species in gene trees for PhylteR. The effect was larger when more outliers were removed (Figure 7, right), the gain in gCF reaching more than 6% for some branches with PhylteR outliers removal (max 5% for TreeShrink). We observed that the gain in concordance was higher for branches that initially had a high gCF, and smaller for poorly supported nodes (plain dots versus circles in Figure 7). This may be due to an easier identification of outliers on a ‘clean’ background (many gene trees supporting the same node, leading to high gCF) than on a more noisy one.

**Figure 7.**
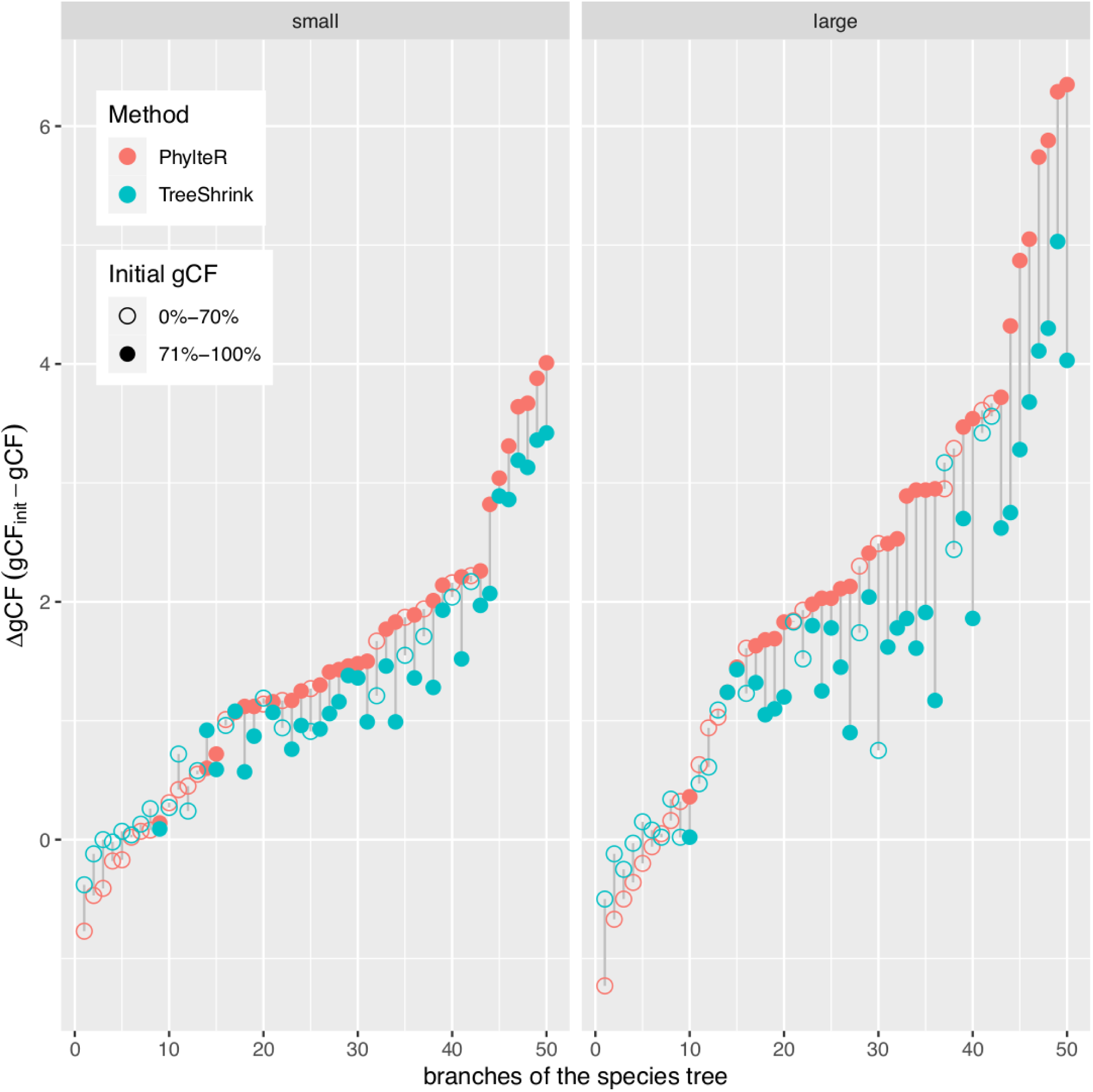
Effect of filtering outliers in gene trees on the gene concordance factor (gCF) of each branch of the Carnivora species tree. The gain in concordance (ΔgCF, y-axis) is plotted for each branch of the species tree (dots), separating PhylteR (pink) and TreeShrink (blue). Branches are ordered by increasing ΔgCF for the PhylteR outliers. The results for the two collections of outliers (small and large) are displayed side by side.

Note that gCF, which captures topological differences between the gene trees and the species tree, exhibits a notable increase but does not attain its maximum value. This observation might be indicative of some of the incongruences between gene and species trees within the Carnivora dataset to be attributed to ILS. These potential ILS-related incongruences appear not to be identified as outliers by PhylteR, as suggested by the results of our simulations (see above).

## Discussion

In phylogenomics, incongruence between gene trees, resulting from a myriad of possible technical and analytical issues, or from biological processes, is known to lead to errors in species tree inference (Philippe et al. 2017). A common practice in phylogenomics thus consists of scanning individual gene trees by eye, trying to spot species or group of species weirdly placed in gene trees, suspicious long branches, apparent groups of paralogues, etc. and discarding them prior to the concatenation of the genes (supermatrix approach) or to the assembly of the gene trees into a species tree (supertree and coalescent-based approaches). This hard work is not only time-consuming and laborious, it is also questionable: what is the objectivity in this practice? Is the eye (and the brain) capable of looking at tens of thousands of gene trees at the same time? How reproducible is such a practice? Etc.

Here, with PhylteR, we propose a way of analysing large collections of gene trees by using an automatic method that can simultaneously analyse a large collection of distance matrices (retrieved from gene trees), identify the common signal between these matrices, and identify elements (outliers) in some of these matrices that are responsible for a decrease in concordance. By using a process where these outliers are automatically and iteratively removed, we propose a new way of efficiently identifying them.

Evaluating a method for its capacity to accurately identify errors in phylogenomics datasets is a difficult task. As for any inference method, we use simulations. However, simulating the processes that result in errors (in our case, outliers in phylogenomics data) has no standard solution: sources of errors are numerous, they combine with each other through all phylogenomic pipelines, sometimes with unpredictable results. So we restricted ourselves to simulating a feature intrinsically detectable by PhylteR, that is, changes in the phylogenetic placements of some species in some gene trees. Further evaluation would involve an independent simulation pipeline, not informed by the hypothesis behind the inference method (Biller et al. 2016), which is by definition outside the scope of the description of the inference method. The simple simulations we performed revealed that outliers corresponding to misplacement of species in a few gene trees was easy to detect with PhylteR but not with TreeShrink. However, this is not surprising *a posteriori*: TreeShrink (Mai and Mirarab 2018) is designed to detect abnormally long branches in collection of gene trees, while we considered here as outliers species that changed position in some gene trees because of horizontal gene transfers; these outliers are not necessarily associated with longer branches.

A better way to examine the advantages of a method over another is to explore biological data. To this end, we evaluated PhylteR and compared it with TreeShrink by looking at some properties associated with gene sequences, and testing possible enrichment of these properties in the list of detected outliers. We observed an enrichment of short sequences, which was anticipated (short sequences carry less phylogenetic signal) and confirmed previous results (Shen et al. 2016).

A notable difference that we observed between PhylteR and TreeShrink, confirming the results obtained on the simple simulated examples, is the duplication score computed here: outliers identified with PhylteR seemed to be highly enriched in gene sequences having experienced more duplications, according to the reconciliation analysis performed. Note, however, that we need to be cautious with this measure: being based solely on a topological comparison between gene and species trees, it cannot distinguish between true paralogy, and other processes (biological or artefactual) leading to a species in a gene tree to have a position that is not concordant with its position in the other gene trees. Horizontal gene transfers (HGT) for instance, may lead to high duplication scores according to our approach when none occurred (even though HGT is thought to be anecdotal in the carnivora dataset). Similarly, artefactual reasons such as long branch attraction, annotation error or alignment error can lead to misplacements of species in some gene trees.

A more direct way of testing the ability of PhylteR to detect hidden paralogous sequences was to focus on a specific gene family known to be extremely diverse because of multiple duplication events, the KZNF family (Huntley et al. 2006; Liu et al. 2014). We observed a clear enrichment of sequences belonging to this peculiar family in the list of outlier sequences identified by PhylteR, as compared to those identified by TreeShrink or randomly sampled. This capacity of PhylteR to identify putative paralogs is an important feature, as it was shown earlier that non-orthologous sequences in phylogenomic datasets could have drastic impact on results (Philippe et al. 2017), leading for instance to erroneous branching with high support in the reconstructed species tree in some cases (Philippe et al. 2011).

A final test that we used to validate PhylteR consisted in exploring the syntenic nature (and lack thereof) of the sequences identified as outliers when comparing the species in a pairwise manner. We observed that outlier sequences were often (much more than expected by chance) syntenic-outliers, i.e. sequences associated with a loss of synteny when comparing the two genomes. This provides two kinds of information: on one side, that the “syntenic outliers” and the “phylogenetic outliers” largely overlap, which proves with an argument orthogonal to all the previous ones, that PhylteR (and TreeShrink to a lesser extent) captures an information about erroneous annotations; on the other side, it suggests that many “syntenic outliers” are due to errors and not to biological processes. “Syntenic outliers” are often filtered out before performing rearrangement analyses, because their position is believed to be artefactual (Lucas and Crollius 2017). However sometimes this outlier position is modelled as the result of a biological process (Dalevi and Eriksen 2008). Our analysis supports this artifactual origin in Carnivora, though some syntenic outliers might originate from retrotranscription or translocations.

Incomplete Lineage Sorting (ILS) is a known source of incongruence among gene trees and between gene trees and the species tree. This biological process, where ancestral polymorphism is maintained across various speciation events, leads to different portions of the genomes having different evolutionary histories. With simulations we could vary the level of Incomplete Lineage Sorting in the datasets and evaluate the impact it had on the ability of PhyleR (and TreeShrink) to correctly identify outliers. For both methods, we saw no effect of increasing the level of ILS on the precision and sensitivity values. For TreeShrink, it is hard to conclude anything, because the initial values were very low (close to 0) so that any negative effect would have been undetectable. For PhylteR however, where precision and sensitivity were high, this absence of effect reveals that PhylteR does not detect sequences that have experienced ILS as outliers. Whether this is positive or negative can be discussed. On the one hand, it was shown earlier that ILS-related incongruences among gene trees could have a detrimental effect for species tree reconstruction with supermatrix approaches (Degnan and Rosenberg 2006). On the other hand, ILS-induced incongruence is a true biological signal that many species tree inference methods, namely the coalescence-based ones, can now handle (Liu et al. 2010; Zhang et al. 2018). In this context, getting rid of these incongruences may be seen as detrimental, because it removes a meaningful signal that can be accommodated by these methods. However, the real contribution of ILS to gene tree incongruences is something that is rarely measured, but in mammals for instance, it was shown to be rare (Scornavacca and Galtier 2017). We can advance two reasons why PhylteR is not sensitive to ILS. First, ILS preferentially affects short branches of the species tree, i.e. speciation events separated by a short amount of time, which leads to a limited effect in the pairwise distance matrices manipulated by PhylteR. Second, when ILS changes the branching pattern of three clades (or species), it is expected that around 50% of each alternative topology to the true one is observed across all gene trees. When the “compromise” matrix is built in the PhylteR pipeline, this signal will thus likely be averaged out.

Here we focused on the identification of outliers in collection of gene trees in order to remove them prior to phylogenetic inference with supermatrix, supertree or coalescent-based methods. But other usage of the tool we present here can be anticipated. First, because the PhylteR method consists of comparing matrices (in this case phylogenetic distance matrices), it is easy to imagine applying the method without computing gene trees, directly on matrices extracted from multiple sequence alignments (MSA), one matrix per gene. In this sense, comparing PhylteR with MSA-based filtering tools could be a worthwhile follow-up of this work. Second, correctly identifying and removing outliers from phylogenomic datasets could be of interest beyond species tree reconstruction. For instance, it appears to be crucial when using statistical methods based on the ratio of nonsynonymous over synonymous substitution rates (*d_N_/d_S_* ratio) to detect adaptive molecular evolution (see Yang and Bielawski 2000 for a review), or for correctly inferring ancestral sequences (Yang et al. 1995) from sequences of extant species. Finally, using a tool like PhylteR is not only useful for cleaning the data. The in-depth exploration of the outliers detected and the study of the reasons why they were detected as such can give important insights into the evolutionary history of these sequences, for instance allowing for the identification of horizontally transferred or duplicated genes.

## Conclusion

We created PhylteR, a tool to explore phylogenomics dataset and detect outlier gene sequences. Instead of fully removing rogue taxa or full outlier gene family, PhylteR precisely identifies what sequences in what gene family should be removed to increase concordance between genes. In doing so it accurately spots gene sequences with low phylogenetic signal, genes with saturated signal leading to long branches, paralogous genes, genes associated with synteny breaks and other sequences that are dubious in gene phylogenies for any possible reason.

## Supporting information

supp mat

## Acknowledgments

Work was funded by ANR Grant 18-CE02-0007 (Sthoriz) to DMDV, ANR Grant 19-CE45-0010 (Evoluthon) to ET, and European Research Council grant ERC-2015-CoG-683257 (ConvergeAnt project) to FD. This is contribution ISEM 2023-XXX of the Institut des Sciences de l’Evolution de Montpellier. We thank Mélodie Bastian for help with the simulations and two anonymous reviewers for helpful comments.

## Software availability

PhylteR is a package written in R language (R Core Team 2023) available on CRAN (https://cran.r-project.org/web/packages/phylter/index.html) for the latest stable version and on GitHub (https://github.com/damiendevienne/phylter) for the latest development version. The latest version of PhylteR is also distributed as a Singularity container (https://cloud.sylabs.io/library/theo.treecou/tool/phylter_singularity) and a docker container (https://hub.docker.com/r/treecoutheo/phylter_docker). Extensive documentation can be found at https://damiendevienne.github.io/phylter/index.html.

## Data Availability

The documented code of PhylteR is available at https://github.com/damiendevienne/phylter along with a thorough documentation. All data and scripts used in this study are available on the dedicated GitHub repository available at https://github.com/damiendevienne/phylter-data/.

