## Supplementary material for "PhylteR: efficient identification of outlier sequences in phylogenomic datasets": supp mat

### Supplementary methods

For simulating the gene trees using Simphy, we used the following command, replacing SPECIESTREENEWICK and ILSLEVEL by their values, given just after:

```
simphy_lnx64 -rs 1 -rl f:500 -rg 1 -s SPECIESTREENEWICK -sg
f:7.62125 -sp ILSLEVEL -su f:2.2e-09 -ld f:0 -lb f:0 -lt f:1e-10
-lg f:0 -o simphysimul -om 1 -ot 0 -od 1 -op 1 -oc 1 -lk 0 -hg
f:100 -ol 1
```

**SPECIESTREENEWICK**:

```
'(Manis_javanica:563972501,(((Paradoxurus_hermaphroditus:178299388
,((Cryptoprocta_ferox:113225468,(Suricata_suricatta:42276894,(Helo
gale_parvula:35627804,Mungos_mungo:35627804):6649090):70948574):27
721304,((Hyaena_hyaena:21169725,Crocuta_Crocuta:21169725):5080201,
(Proteles_cristatus:4459256,Proteles_septentrionalis:4459256):2179
0670):114696846):37352616):15821653,((Neofelis_nebulosa:35249192,(
(Panthera_leo:14904805,Panthera_onca:14904805):13788677,(Panthera_
pardus:20008477,Panthera_tigris:20008477):8685005):6555710):154610
15,(Prionailurus_bengalensis:41904186,(Felis_catus:36061165,((Lynx
_pardinus:13846455,Lynx_canadensis:13846455):19799374,(Acinonyx_ju
batus:24562910,Puma_concolor:24562910):9082919):2415336):5843021):
8806021):143410834):111178330,(((Canis_familiaris:20610593,Lycaon_
pictus:20610593):16674959,(Otocyon_megalotis:33511470,Vulpes_vulpe
s:33511470):3774082):219916194,((Ailuropoda_melanoleuca:76527633,(
Ursus_maritimus:30159998,(Ursus_thibetanus:16629885,(Ursus_america
nus:13861784,Ursus_arctos:13861784):2768101):13530113):46367635):1
35019564,(((Phoca_vitulina:66480327,(Neomonachus_schauinslandi:428
62823,(Leptonychotes_weddellii:38322355,Mirounga_angustirostris:38
322355):4540468):23617504):46423630,(Odobenus_rosmarus:60799123,(C
allorhinus_ursinus:33538848,(Arctocephalus_gazella:29177543,(Eumet
opias_jubatus:17584216,Zalophus_californianus:17584216):11593327):
4361305):27260275):52104834):89655108,(Spilogale_gracilis:16765364
7,(Ailurus_fulgens:155264004,((Potos_flavus:108127883,(Nasua_naric
a:86590883,(Bassariscus_sumichrasti:42191209,Procyon_lotor:4219120
9):44399674):21537000):35853476,(Taxidea_taxus:84460064,(Mellivora
_capensis:70679485,(Gulo_gulo:65876915,((Mustela_putorius:34170509
,Neovison_vison:34170509):20370272,(Pteronura_brasiliensis:4305120
2,(Enhydra_lutris:26877386,Lutra_lutra:26877386):16173816):1148957
9):11336134):4802570):13780579):59521295):11282645):12389643):3490
5418):8988132):45654549):48097625):258673130);'
```

**ILSLEVEL**:

`10:` (NO-ILS)

`100000:` (LOW-ILS)

`200000:` (MODERATE-ILS)

`500000:` (HIGH-ILS)

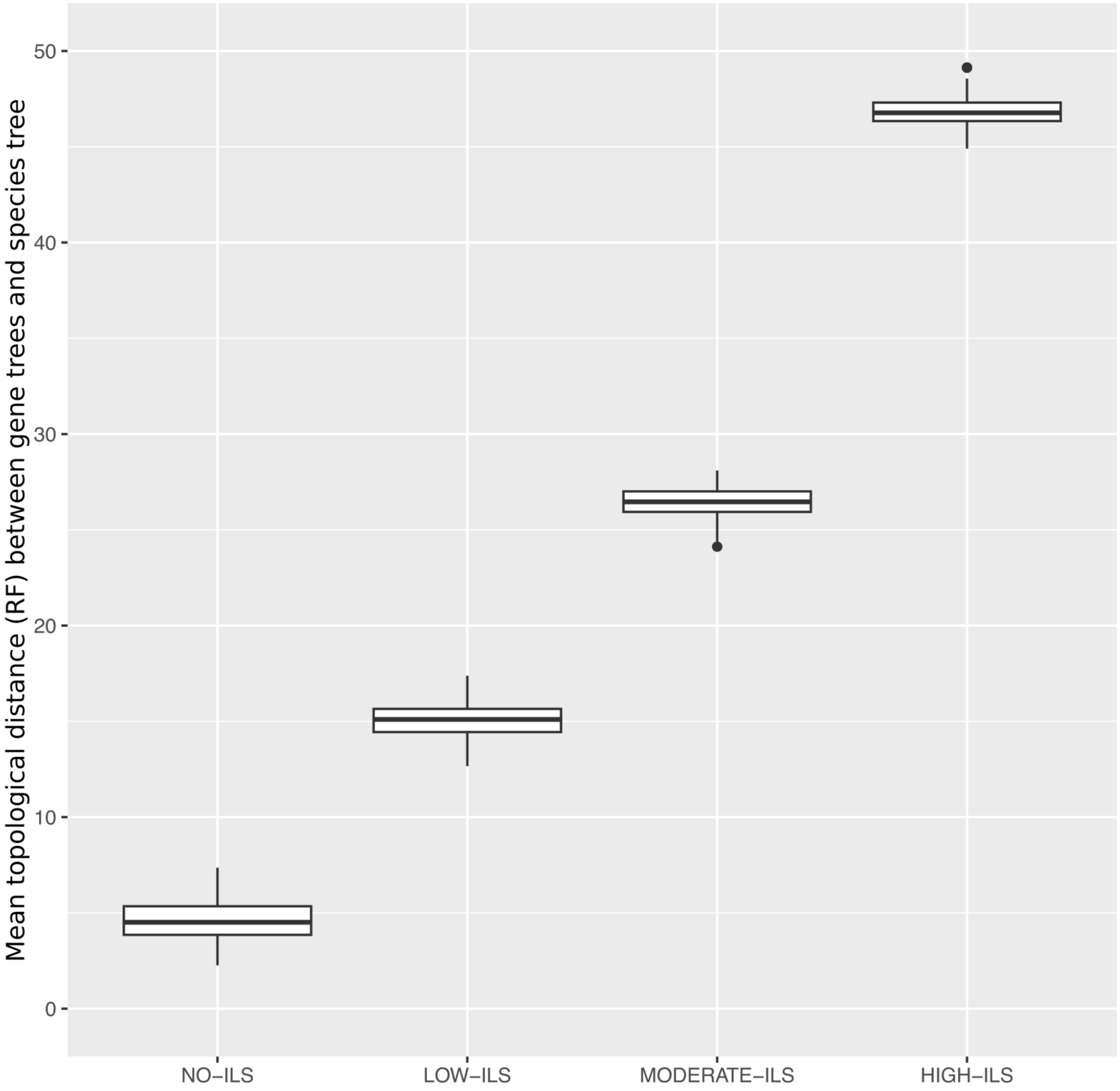


**Figure S1. Impact of Incomplete Lineage Sorting on the mean RF distance between the simulated gene trees and the species tree.** The mean topological distance observed when NO-ILS is present is imputable to Horizontal Gene Transfers (HGTs) only.

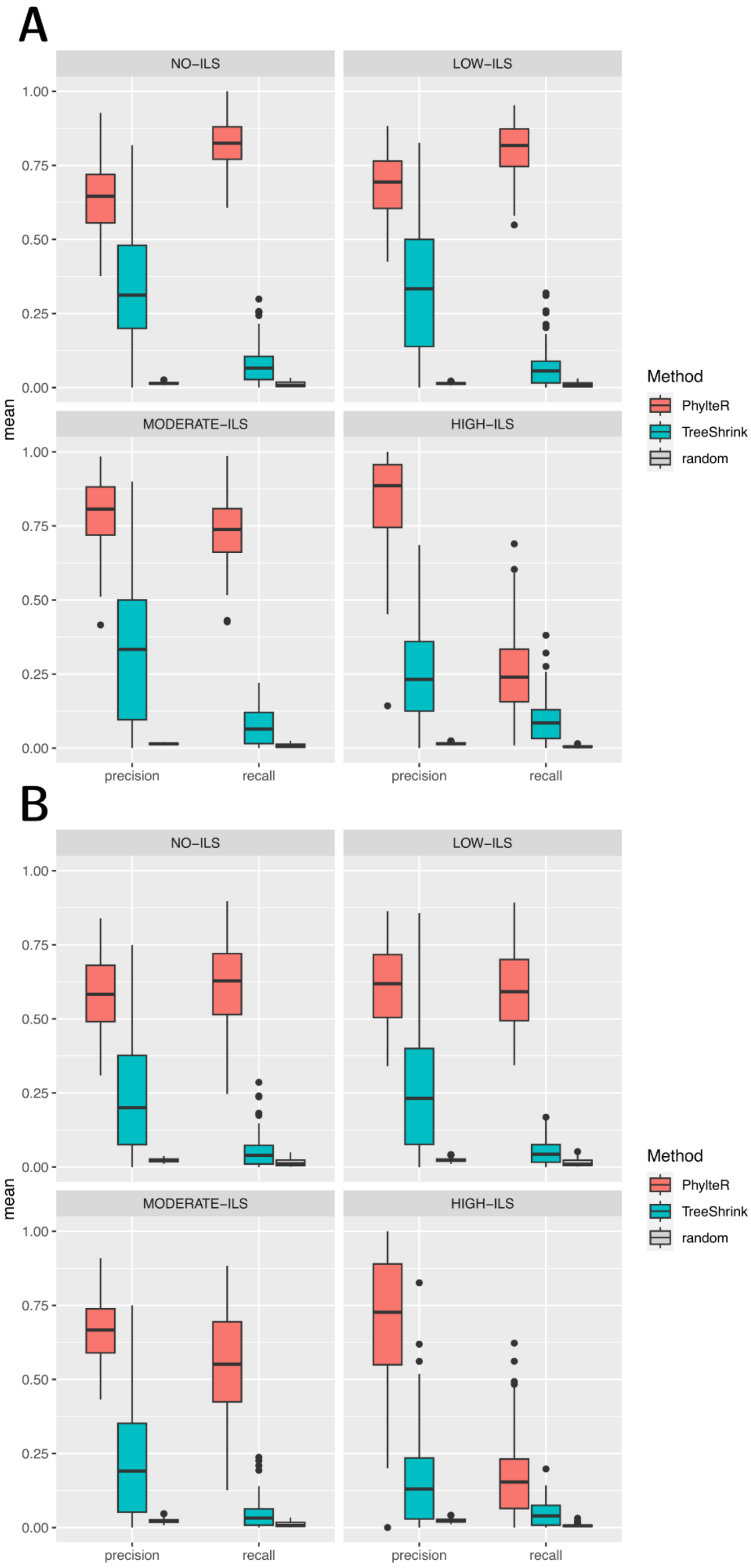


**Figure S2. Comparison of the precision and recall (or sensitivity) of the PhylteR and the TreeShrink outlier detection methods.** Either a maximum of 10 outlier species (**A**) or a (theoretical) maximum of 53 species (**B**) are allowed per gene tree. Each time, the four conditions of Incomplete Lineage Sorting (ILS) are given, one per panel.

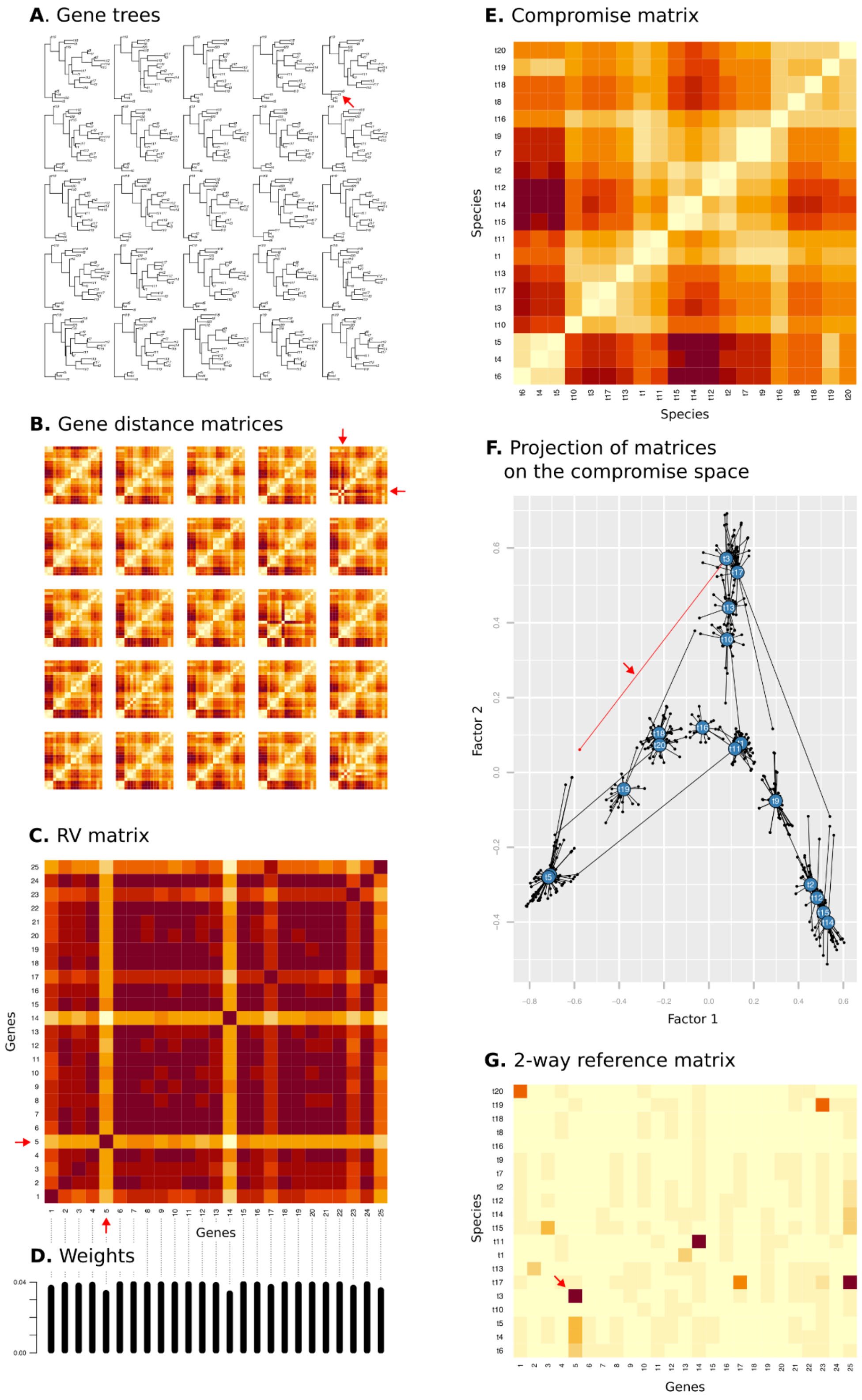


**Figure S3. Illustration of the different steps of the PhylteR process depicted in Figure 1 (main text).** The red arrow identifies on each step, one of the outliers of the dataset, namely species t3 in gene 5.

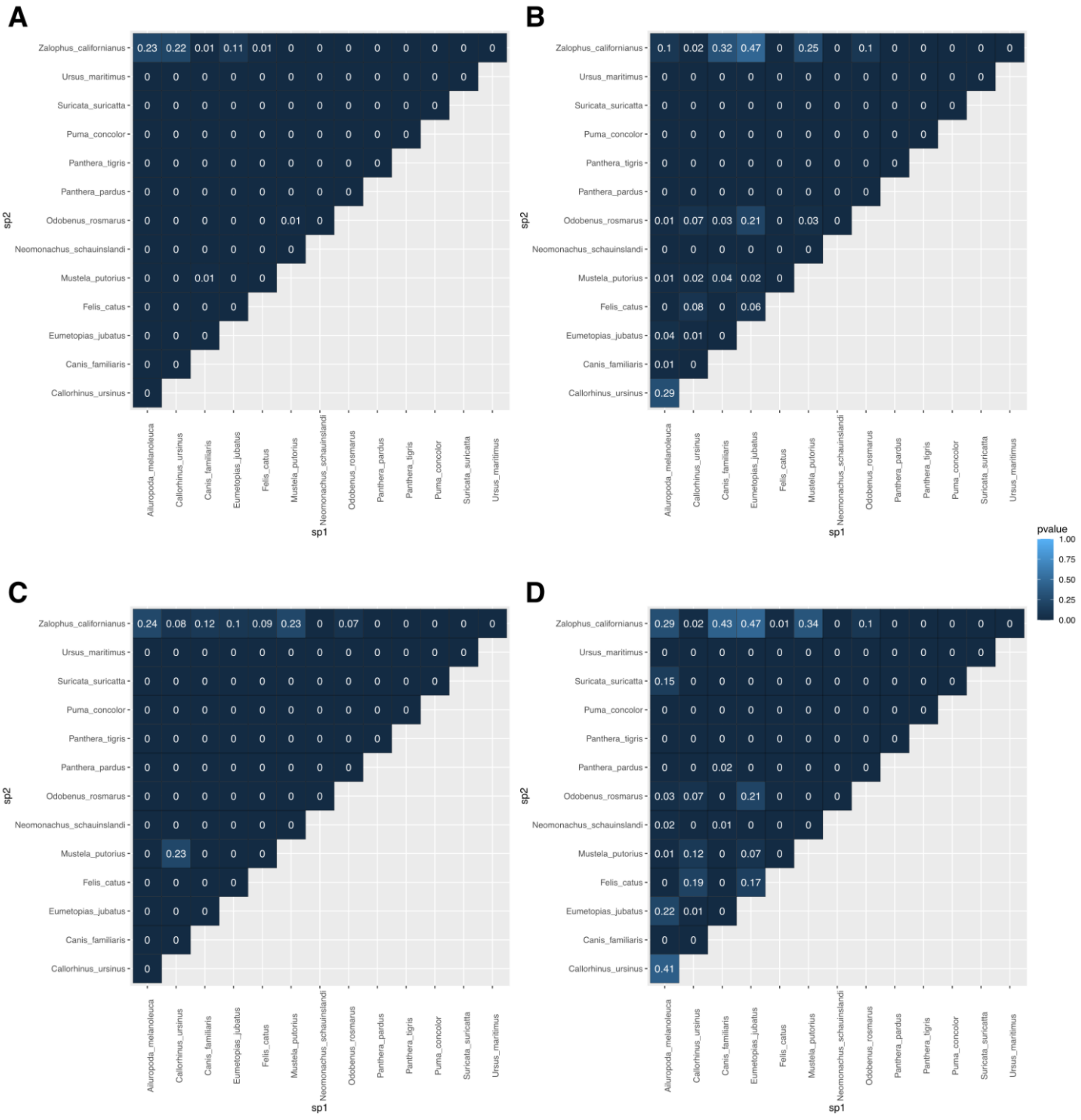


**Figure S4. Analysis of the synteny breaks in the list of outliers.** Each heatmap represents all pairwise comparisons between the 14 species of interest. For each comparison, the p-value associated with the probability of getting the observed number of syntenic outliers in the list of outliers is indicated. The first row (A and B) are the results with PhylteR outliers, the second row (C and D) are for TreeShrink outliers. The two columns represent the two sizes of outlier lists, small (A and C) and large (B and D).

**Table S1. Genomes used for synteny breakage analysis.**

| Species | Family | Accession | AssemblyName | # scaffolds |
|---|---|---|---|---|
| *Ailuropoda melanoleuca* | Ursidae | GCF_000004335.2 | AilMel_1.0 | 1913 |
| *Callorhinus ursinus* | Otariidae | GCF_003265705.1 | ASM326570v1 | 146 |
| *Canis lupus* | Canidae | GCF_000002285.3 | CanFam3.1 | 59 |
| *Eumetopias jubatus* | Otariidae | GCF_004028035.1 | ASM402803v1 | 323 |
| *Felis catus* | Felidae | GCF_000181335.3 | Felis_catus_9.0 | 25 |
| *Mustela putorius* | Mustelidae | GCF_000215625.1 | MusPutFur1.0 | 457 |
| *Neomonachus schauinslandi* | Phocidae | GCF_002201575.1 | ASM220157v1 | 273 |
| *Odobenus rosmarus* | Odobenidae | GCF_000321225.1 | Oros_1.0 | 1170 |
| *Panthera pardus* | Felidae | GCF_001857705.1 | PanPar1.0 | 289 |
| *Panthera tigris* | Felidae | GCF_000464555.1 | PanTig1.0 | 505 |
| *Puma concolor* | Felidae | GCF_003327715.1 | PumCon1.0 | 48 |
| *Suricata suricatta* | Herpestidae | GCF_006229205.1 | meerkat_22Aug2017_6uvM2_HiC | 25 |
| *Ursus maritimus* | Ursidae | GCF_000687225.1 | UrsMar_1.0 | 314 |
| *Zalophus californianus* | Otariidae | GCF_900631625.1 | zalCal2.2 | 27 |

**Table S2. Comparisons of genomes for the synteny analysis with PhylteR outlier lists.**

| | | | Phylter Outliers (small list) | | | Phylter outliers (large list) | | |
|---|---|---|---|---|---|---|---|---|
| **Species pair** | **Genes** | **Syntenic outliers** | **Total** | **syntenic** | **P-value** | **Total** | **Syntenic** | **P-value** |
| *Ailuropoda melanoleuca - Callorhinus ursinus* | 6493 | 558 | 56 | 14 | 1.986E-04 | 182 | 29 | 7.696E-04 |
| *Ailuropoda melanoleuca - Canis familiaris* | 12192 | 753 | 247 | 33 | 2.215E-05 | 740 | 66 | 1.484E-03 |
| *Ailuropoda melanoleuca - Eumetopias jubatus* | 7543 | 682 | 68 | 17 | 8.122E-05 | 223 | 36 | 3.977E-04 |
| *Ailuropoda melanoleuca - Felis catus* | 11229 | 656 | 144 | 19 | 7.040E-04 | 450 | 46 | 1.409E-04 |
| *Ailuropoda melanoleuca - Mustela putorius* | 11720 | 770 | 116 | 17 | 1.458E-03 | 445 | 56 | 1.740E-06 |
| *Ailuropoda melanoleuca - Neomonachus schauinslandi* | 12335 | 797 | 142 | 29 | 2.358E-08 | 454 | 53 | 1.854E-05 |
| *Ailuropoda melanoleuca - Odobenus rosmarus* | 12214 | 913 | 147 | 26 | 3.071E-05 | 460 | 56 | 1.847E-04 |
| *Ailuropoda melanoleuca - Panthera pardus* | 12123 | 733 | 150 | 22 | 9.565E-05 | 514 | 52 | 1.682E-04 |
| *Ailuropoda melanoleuca - Panthera tigris* | 11725 | 840 | 165 | 66 | 3.106E-33 | 483 | 99 | 1.735E-22 |
| *Ailuropoda melanoleuca - Puma concolor* | 9907 | 894 | 119 | 49 | 3.124E-21 | 364 | 93 | 2.337E-21 |
| *Ailuropoda melanoleuca - Suricata suricatta* | 10368 | 730 | 126 | 70 | 6.374E-48 | 378 | 92 | 2.302E-27 |
| *Ailuropoda melanoleuca - Ursus maritimus* | 11924 | 771 | 145 | 54 | 9.758E-28 | 400 | 78 | 3.807E-19 |
| *Ailuropoda melanoleuca - Zalophus californianus* | 5506 | 586 | 47 | 7 | 2.288E-01 | 159 | 20 | 2.450E-01 |
| *Callorhinus ursinus - Canis familiaris* | 6979 | 146 | 143 | 13 | 7.985E-06 | 409 | 18 | 2.083E-03 |

| | | | Phylter Outliers (small list) | | | Phylter outliers (large list) | | |
|---|---|---|---|---|---|---|---|---|
| **Species pair** | **Genes** | **Syntenic outliers** | **Total** | **syntenic** | **P-value** | **Total** | **Syntenic** | **P-value** |
| *Callorhinus ursinus - Eumetopias jubatus* | 6660 | 105 | 40 | 5 | 3.756E-04 | 113 | 9 | 6.521E-05 |
| *Callorhinus ursinus - Felis catus* | 6309 | 92 | 72 | 8 | 8.385E-06 | 205 | 10 | 7.430E-04 |
| *Callorhinus ursinus - Mustela putorius* | 6616 | 182 | 50 | 7 | 3.901E-04 | 182 | 7 | 2.346E-01 |
| *Callorhinus ursinus - Neomonachus schauinslandi* | 7147 | 186 | 73 | 15 | 3.757E-10 | 207 | 18 | 6.373E-06 |
| *Callorhinus ursinus - Odobenus rosmarus* | 7061 | 355 | 75 | 11 | 1.199E-03 | 207 | 22 | 6.725E-04 |
| *Callorhinus ursinus - Panthera pardus* | 7003 | 161 | 86 | 11 | 3.722E-06 | 261 | 22 | 1.039E-07 |
| *Callorhinus ursinus - Panthera tigris* | 6617 | 212 | 82 | 40 | 5.197E-39 | 222 | 53 | 6.342E-33 |
| *Callorhinus ursinus - Puma concolor* | 6086 | 246 | 58 | 31 | 9.747E-29 | 173 | 46 | 3.465E-26 |
| *Callorhinus ursinus - Suricata suricatta* | 5953 | 139 | 58 | 40 | 4.026E-54 | 165 | 46 | 6.236E-39 |
| *Callorhinus ursinus - Ursus maritimus* | 6760 | 160 | 68 | 29 | 3.304E-30 | 165 | 35 | 1.819E-24 |
| *Callorhinus ursinus - Zalophus californianus* | 4782 | 71 | 17 | 1 | 2.249E-01 | 70 | 3 | 8.492E-02 |
| *Canis familiaris - Eumetopias jubatus* | 8091 | 190 | 182 | 17 | 1.134E-06 | 485 | 26 | 6.111E-05 |
| *Canis familiaris - Felis catus* | 11863 | 216 | 254 | 16 | 1.623E-05 | 734 | 28 | 1.584E-04 |
| *Canis familiaris - Mustela putorius* | 12383 | 301 | 239 | 13 | 5.692E-03 | 728 | 36 | 3.754E-05 |
| *Canis familiaris - Neomonachus schauinslandi* | 13119 | 358 | 271 | 29 | 3.072E-10 | 808 | 48 | 2.678E-07 |
| *Canis familiaris - Odobenus rosmarus* | 12955 | 517 | 271 | 29 | 1.299E-06 | 806 | 65 | 3.656E-08 |

| | | | Phylter Outliers (small list) | | | Phylter outliers (large list) | | |
|---|---|---|---|---|---|---|---|---|
| **Species pair** | **Genes** | **Syntenic outliers** | **Total** | **syntenic** | **P-value** | **Total** | **Syntenic** | **P-value** |
| *Canis familiaris - Panthera pardus* | 12919 | 313 | 287 | 24 | 1.293E-07 | 849 | 40 | 3.868E-05 |
| *Canis familiaris - Panthera tigris* | 12405 | 407 | 286 | 75 | 2.215E-47 | 795 | 108 | 2.913E-39 |
| *Canis familiaris - Puma concolor* | 10525 | 444 | 217 | 53 | 1.990E-26 | 631 | 87 | 7.318E-24 |
| *Canis familiaris - Suricata suricatta* | 10948 | 276 | 244 | 83 | 8.032E-74 | 674 | 100 | 6.369E-52 |
| *Canis familiaris - Ursus maritimus* | 12614 | 355 | 276 | 61 | 9.473E-38 | 746 | 83 | 1.835E-28 |
| *Canis familiaris - Zalophus californianus* | 5918 | 106 | 119 | 7 | 5.254E-03 | 367 | 10 | 1.201E-01 |
| *Eumetopias jubatus - Felis catus* | 7318 | 132 | 77 | 9 | 8.797E-06 | 231 | 13 | 2.539E-04 |
| *Eumetopias jubatus - Mustela putorius* | 7666 | 251 | 69 | 12 | 1.957E-06 | 226 | 16 | 2.972E-03 |
| *Eumetopias jubatus - Neomonachus schauinslandi* | 8287 | 241 | 95 | 27 | 5.158E-20 | 247 | 29 | 1.003E-10 |
| *Eumetopias jubatus - Odobenus rosmarus* | 8178 | 422 | 89 | 19 | 9.234E-08 | 236 | 27 | 7.787E-05 |
| *Eumetopias jubatus - Panthera pardus* | 8109 | 205 | 107 | 18 | 1.354E-10 | 312 | 28 | 3.966E-09 |
| *Eumetopias jubatus - Panthera tigris* | 7685 | 273 | 99 | 43 | 6.526E-37 | 258 | 55 | 1.328E-28 |
| *Eumetopias jubatus - Puma concolor* | 7099 | 293 | 78 | 36 | 8.046E-30 | 219 | 48 | 1.202E-22 |
| *Eumetopias jubatus - Suricata suricatta* | 6905 | 197 | 72 | 48 | 7.701E-59 | 212 | 56 | 2.380E-40 |
| *Eumetopias jubatus - Ursus maritimus* | 7853 | 211 | 102 | 42 | 3.399E-40 | 236 | 50 | 7.035E-32 |

| | | | Phylter Outliers (small list) | | | Phylter outliers (large list) | | |
|---|---|---|---|---|---|---|---|---|
| **Species pair** | **Genes** | **Syntenic outliers** | **Total** | **syntenic** | **P-value** | **Total** | **Syntenic** | **P-value** |
| *Eumetopias jubatus - Zalophus californianus* | 5390 | 91 | 33 | 2 | 1.063E-01 | 104 | 4 | 9.814E-02 |
| *Felis catus - Mustela putorius* | 11436 | 235 | 143 | 12 | 3.785E-05 | 459 | 24 | 2.488E-05 |
| *Felis catus - Neomonachus schauinslandi* | 12019 | 286 | 165 | 18 | 7.406E-08 | 493 | 29 | 6.411E-06 |
| *Felis catus - Odobenus rosmarus* | 11890 | 443 | 153 | 17 | 5.489E-05 | 474 | 40 | 1.085E-06 |
| *Felis catus - Panthera pardus* | 11887 | 176 | 149 | 14 | 4.142E-08 | 408 | 23 | 3.412E-08 |
| *Felis catus - Panthera tigris* | 11482 | 258 | 161 | 41 | 3.468E-32 | 396 | 55 | 1.303E-28 |
| *Felis catus - Puma concolor* | 9687 | 303 | 115 | 30 | 6.281E-20 | 306 | 53 | 2.375E-25 |
| *Felis catus - Suricata suricatta* | 10140 | 210 | 121 | 56 | 9.431E-64 | 351 | 67 | 2.597E-47 |
| *Felis catus - Ursus maritimus* | 11628 | 227 | 161 | 39 | 3.804E-32 | 448 | 48 | 1.002E-22 |
| *Felis catus - Zalophus californianus* | 5268 | 72 | 57 | 4 | 7.342E-03 | 177 | 5 | 9.343E-02 |
| *Mustela putorius - Neomonachus schauinslandi* | 12565 | 389 | 141 | 17 | 1.647E-06 | 474 | 33 | 1.164E-05 |
| *Mustela putorius - Odobenus rosmarus* | 12441 | 522 | 139 | 13 | 5.622E-03 | 475 | 33 | 3.103E-03 |
| *Mustela putorius - Panthera pardus* | 12334 | 323 | 153 | 17 | 5.161E-07 | 523 | 34 | 9.247E-07 |
| *Mustela putorius - Panthera tigris* | 11905 | 415 | 154 | 58 | 3.658E-45 | 494 | 81 | 4.598E-33 |
| *Mustela putorius - Puma concolor* | 10064 | 428 | 108 | 38 | 2.150E-25 | 370 | 62 | 2.780E-21 |
| *Mustela putorius - Suricata suricatta* | 10496 | 303 | 115 | 65 | 2.327E-71 | 372 | 78 | 1.550E-46 |

| | | | Phylter Outliers (small list) | | | Phylter outliers (large list) | | |
|---|---|---|---|---|---|---|---|---|
| **Species pair** | **Genes** | **Syntenic outliers** | **Total** | **syntenic** | **P-value** | **Total** | **Syntenic** | **P-value** |
| *Mustela putorius - Ursus maritimus* | 12097 | 327 | 148 | 44 | 4.341E-34 | 447 | 62 | 2.157E-27 |
| *Mustela putorius - Zalophus californianus* | 5537 | 174 | 37 | 6 | 9.131E-04 | 159 | 7 | 2.327E-01 |
| *Neomonachus schauinslandi - Odobenus rosmarus* | 13241 | 568 | 155 | 29 | 1.555E-11 | 462 | 46 | 8.479E-08 |
| *Neomonachus schauinslandi - Panthera pardus* | 13160 | 357 | 190 | 28 | 2.287E-13 | 586 | 46 | 7.056E-11 |
| *Neomonachus schauinslandi - Panthera tigris* | 12635 | 413 | 188 | 65 | 1.386E-49 | 539 | 88 | 5.334E-38 |
| *Neomonachus schauinslandi - Puma concolor* | 10711 | 449 | 146 | 56 | 9.144E-40 | 421 | 83 | 3.797E-34 |
| *Neomonachus schauinslandi - Suricata suricatta* | 11126 | 328 | 134 | 72 | 4.327E-76 | 420 | 85 | 1.469E-48 |
| *Neomonachus schauinslandi - Ursus maritimus* | 12855 | 380 | 172 | 57 | 9.094E-45 | 473 | 66 | 7.579E-27 |
| *Neomonachus schauinslandi - Zalophus californianus* | 6052 | 163 | 54 | 15 | 5.205E-12 | 183 | 18 | 1.637E-06 |
| *Odobenus rosmarus - Panthera pardus* | 12988 | 538 | 181 | 24 | 4.466E-07 | 583 | 47 | 9.022E-06 |
| *Odobenus rosmarus - Panthera tigris* | 12447 | 613 | 189 | 72 | 1.344E-45 | 540 | 100 | 3.563E-32 |
| *Odobenus rosmarus - Puma concolor* | 10558 | 665 | 142 | 56 | 6.060E-31 | 415 | 93 | 1.767E-28 |
| *Odobenus rosmarus - Suricata suricatta* | 11012 | 521 | 137 | 69 | 2.882E-55 | 421 | 91 | 1.960E-36 |
| *Odobenus rosmarus - Ursus maritimus* | 12672 | 554 | 172 | 54 | 5.486E-32 | 479 | 72 | 9.482E-21 |
| *Odobenus rosmarus - Zalophus californianus* | 5973 | 351 | 57 | 10 | 1.561E-03 | 183 | 16 | 7.072E-02 |

| | | | Phylter Outliers (small list) | | | Phylter outliers (large list) | | |
|---|---|---|---|---|---|---|---|---|
| **Species pair** | **Genes** | **Syntenic outliers** | **Total** | **syntenic** | **P-value** | **Total** | **Syntenic** | **P-value** |
| *Panthera pardus - Panthera tigris* | 12541 | 381 | 183 | 55 | 5.716E-40 | 480 | 79 | 1.201E-36 |
| *Panthera pardus - Puma concolor* | 10607 | 428 | 158 | 49 | 1.305E-30 | 395 | 78 | 2.561E-33 |
| *Panthera pardus - Suricata suricatta* | 10955 | 290 | 141 | 69 | 1.152E-72 | 403 | 83 | 3.834E-52 |
| *Panthera pardus - Ursus maritimus* | 12660 | 323 | 189 | 49 | 6.084E-36 | 541 | 62 | 3.828E-24 |
| *Panthera pardus - Zalophus californianus* | 5938 | 135 | 70 | 8 | 1.671E-04 | 230 | 15 | 2.045E-04 |
| *Panthera tigris - Puma concolor* | 10394 | 439 | 119 | 44 | 1.932E-30 | 334 | 69 | 1.787E-29 |
| *Panthera tigris - Suricata suricatta* | 10583 | 331 | 141 | 83 | 1.168E-90 | 385 | 105 | 1.344E-72 |
| *Panthera tigris - Ursus maritimus* | 12504 | 363 | 165 | 51 | 8.915E-39 | 477 | 74 | 6.926E-34 |
| *Panthera tigris - Zalophus californianus* | 5593 | 203 | 67 | 41 | 1.732E-43 | 200 | 48 | 1.454E-27 |
| *Puma concolor - Suricata suricatta* | 9020 | 359 | 120 | 78 | 1.140E-81 | 322 | 99 | 1.704E-63 |
| *Puma concolor - Ursus maritimus* | 10452 | 413 | 128 | 49 | 4.479E-36 | 365 | 72 | 2.534E-31 |
| *Puma concolor - Zalophus californianus* | 5164 | 209 | 44 | 24 | 8.684E-23 | 161 | 37 | 6.387E-19 |
| *Suricata suricatta - Ursus maritimus* | 10741 | 270 | 145 | 83 | 3.697E-98 | 394 | 93 | 3.550E-67 |
| *Suricata suricatta - Zalophus californianus* | 5123 | 132 | 47 | 33 | 1.339E-43 | 149 | 42 | 4.734E-34 |
| *Ursus maritimus - Zalophus californianus* | 5717 | 158 | 69 | 37 | 5.993E-41 | 171 | 42 | 1.287E-29 |

**Table S3. Comparisons of genomes for the synteny analysis with TreeShrink outlier lists.**

| | | | TreeShrink Outliers (small list) | | | TreeShrink outliers (large list) | | |
|---|---|---|---|---|---|---|---|---|
| **Species pair** | **Genes** | **Syntenic outliers** | **Total** | **Syntenic outliers** | **P-value** | **Total** | **Syntenic outliers** | **P-value** |
| *Ailuropoda melanoleuca - Canis familiaris* | 12192 | 753 | 215 | 22 | 1.362E-02 | 803 | 71 | 1.248E-03 |
| *Ailuropoda melanoleuca - Eumetopias jubatus* | 7543 | 682 | 111 | 16 | 4.073E-02 | 225 | 24 | 2.240E-01 |
| *Ailuropoda melanoleuca - Felis catus* | 11229 | 656 | 144 | 22 | 3.013E-05 | 351 | 34 | 2.526E-03 |
| *Ailuropoda melanoleuca - Mustela putorius* | 11720 | 770 | 204 | 22 | 1.486E-02 | 781 | 67 | 1.382E-02 |
| *Ailuropoda melanoleuca - Neomonachus schauinslandi* | 12335 | 797 | 168 | 24 | 1.990E-04 | 332 | 32 | 1.509E-02 |
| *Ailuropoda melanoleuca - Odobenus rosmarus* | 12214 | 913 | 198 | 25 | 6.764E-03 | 352 | 36 | 3.365E-02 |
| *Ailuropoda melanoleuca - Panthera pardus* | 12123 | 733 | 188 | 27 | 2.352E-05 | 347 | 34 | 3.668E-03 |
| *Ailuropoda melanoleuca - Panthera tigris* | 11725 | 840 | 209 | 75 | 7.103E-34 | 447 | 93 | 1.248E-21 |
| *Ailuropoda melanoleuca - Puma concolor* | 9907 | 894 | 165 | 59 | 1.268E-21 | 367 | 83 | 1.038E-15 |
| *Ailuropoda melanoleuca - Suricata suricatta* | 10368 | 730 | 176 | 29 | 1.382E-05 | 693 | 56 | 1.514E-01 |
| *Ailuropoda melanoleuca - Ursus maritimus* | 11924 | 771 | 173 | 44 | 1.373E-15 | 384 | 56 | 6.694E-09 |
| *Ailuropoda melanoleuca - Zalophus californianus* | 5506 | 586 | 76 | 12 | 1.046E-01 | 164 | 20 | 2.916E-01 |
| *Callorhinus ursinus - Canis familiaris* | 6979 | 146 | 121 | 8 | 3.720E-03 | 404 | 18 | 1.818E-03 |

| | | | TreeShrink Outliers (small list) | | | TreeShrink outliers (large list) | | |
|---|---|---|---|---|---|---|---|---|
| Species pair | Genes | Syntenic outliers | Total | Syntenic outliers | P-value | Total | Syntenic outliers | P-value |
| *Callorhinus ursinus - Eumetopias jubatus* | 6660 | 105 | 47 | 4 | 6.185E-03 | 47 | 4 | 6.185E-03 |
| *Callorhinus ursinus - Felis catus* | 6309 | 92 | 70 | 3 | 8.176E-02 | 103 | 3 | 1.896E-01 |
| *Callorhinus ursinus - Mustela putorius* | 6616 | 182 | 100 | 7 | 1.985E-02 | 358 | 14 | 1.155E-01 |
| *Callorhinus ursinus - Neomonachus schauinslandi* | 7147 | 186 | 79 | 9 | 1.930E-04 | 79 | 9 | 1.930E-04 |
| *Callorhinus ursinus - Odobenus rosmarus* | 7061 | 355 | 72 | 7 | 6.878E-02 | 72 | 7 | 6.878E-02 |
| *Callorhinus ursinus - Panthera pardus* | 7003 | 161 | 95 | 10 | 5.871E-05 | 95 | 10 | 5.871E-05 |
| *Callorhinus ursinus - Panthera tigris* | 6617 | 212 | 122 | 39 | 1.987E-29 | 180 | 42 | 1.483E-25 |
| *Callorhinus ursinus - Puma concolor* | 6086 | 246 | 113 | 41 | 2.366E-29 | 176 | 48 | 7.046E-28 |
| *Callorhinus ursinus - Suricata suricatta* | 5953 | 139 | 101 | 14 | 6.715E-08 | 298 | 20 | 1.611E-05 |
| *Callorhinus ursinus - Ursus maritimus* | 6760 | 160 | 112 | 24 | 4.920E-17 | 169 | 26 | 1.105E-14 |
| *Callorhinus ursinus - Zalophus californianus* | 4782 | 71 | 37 | 3 | 1.699E-02 | 37 | 3 | 1.699E-02 |
| *Canis familiaris - Eumetopias jubatus* | 8091 | 190 | 153 | 11 | 9.194E-04 | 480 | 28 | 6.433E-06 |
| *Canis familiaris - Felis catus* | 11863 | 216 | 174 | 11 | 3.393E-04 | 642 | 28 | 1.506E-05 |
| *Canis familiaris - Mustela putorius* | 12383 | 301 | 251 | 11 | 4.312E-02 | 1117 | 43 | 1.640E-03 |
| *Canis familiaris - Neomonachus schauinslandi* | 13119 | 358 | 214 | 18 | 2.340E-05 | 683 | 30 | 6.916E-03 |

| | | | TreeShrink Outliers (small list) | | | TreeShrink outliers (large list) | | |
|---|---|---|---|---|---|---|---|---|
| **Species pair** | **Genes** | **Syntenic outliers** | **Total** | **Syntenic outliers** | **P-value** | **Total** | **Syntenic outliers** | **P-value** |
| *Canis familiaris - Odobenus rosmarus* | 12955 | 517 | 243 | 16 | 3.445E-02 | 704 | 48 | 1.903E-04 |
| *Canis familiaris - Panthera pardus* | 12919 | 313 | 229 | 16 | 1.465E-04 | 686 | 26 | 1.593E-02 |
| *Canis familiaris - Panthera tigris* | 12405 | 407 | 257 | 71 | 1.649E-46 | 751 | 91 | 7.629E-29 |
| *Canis familiaris - Puma concolor* | 10525 | 444 | 207 | 61 | 1.913E-35 | 664 | 81 | 8.506E-19 |
| *Canis familiaris - Suricata suricatta* | 10948 | 276 | 224 | 26 | 6.989E-11 | 991 | 41 | 1.025E-03 |
| *Canis familiaris - Ursus maritimus* | 12614 | 355 | 264 | 46 | 8.006E-24 | 771 | 65 | 7.198E-16 |
| *Canis familiaris - Zalophus californianus* | 5918 | 106 | 110 | 3 | 3.151E-01 | 348 | 7 | 4.325E-01 |
| *Eumetopias jubatus - Felis catus* | 7318 | 132 | 83 | 4 | 6.249E-02 | 121 | 4 | 1.744E-01 |
| *Eumetopias jubatus - Mustela putorius* | 7666 | 251 | 125 | 9 | 2.119E-02 | 433 | 20 | 7.451E-02 |
| *Eumetopias jubatus - Neomonachus schauinslandi* | 8287 | 241 | 100 | 13 | 5.783E-06 | 100 | 13 | 5.783E-06 |
| *Eumetopias jubatus - Odobenus rosmarus* | 8178 | 422 | 94 | 7 | 2.109E-01 | 94 | 7 | 2.109E-01 |
| *Eumetopias jubatus - Panthera pardus* | 8109 | 205 | 115 | 10 | 6.337E-04 | 115 | 10 | 6.337E-04 |
| *Eumetopias jubatus - Panthera tigris* | 7685 | 273 | 145 | 45 | 5.308E-31 | 209 | 48 | 1.743E-26 |
| *Eumetopias jubatus - Puma concolor* | 7099 | 293 | 126 | 41 | 1.264E-26 | 200 | 48 | 1.691E-24 |
| *Eumetopias jubatus - Suricata suricatta* | 6905 | 197 | 123 | 18 | 9.139E-09 | 367 | 25 | 3.989E-05 |

| | | | TreeShrink Outliers (small list) | | | TreeShrink outliers (large list) | | |
|---|---|---|---|---|---|---|---|---|
| **Species pair** | **Genes** | **Syntenic outliers** | **Total** | **Syntenic outliers** | **P-value** | **Total** | **Syntenic outliers** | **P-value** |
| *Eumetopias jubatus - Ursus maritimus* | 7853 | 211 | 150 | 31 | 1.774E-19 | 216 | 32 | 1.288E-15 |
| *Eumetopias jubatus - Zalophus californianus* | 5390 | 91 | 37 | 1 | 4.685E-01 | 37 | 1 | 4.685E-01 |
| *Felis catus - Mustela putorius* | 11436 | 235 | 186 | 16 | 1.380E-06 | 669 | 33 | 2.382E-06 |
| *Felis catus - Neomonachus schauinslandi* | 12019 | 286 | 142 | 14 | 7.113E-06 | 204 | 16 | 2.941E-05 |
| *Felis catus - Odobenus rosmarus* | 11890 | 443 | 169 | 16 | 5.713E-04 | 230 | 21 | 1.391E-04 |
| *Felis catus - Panthera pardus* | 11887 | 176 | 129 | 14 | 6.433E-09 | 183 | 16 | 1.212E-08 |
| *Felis catus - Panthera tigris* | 11482 | 258 | 146 | 41 | 4.405E-34 | 269 | 44 | 7.307E-26 |
| *Felis catus - Puma concolor* | 9687 | 303 | 123 | 39 | 3.180E-29 | 246 | 45 | 1.416E-22 |
| *Felis catus - Suricata suricatta* | 10140 | 210 | 150 | 22 | 3.894E-13 | 564 | 28 | 1.410E-05 |
| *Felis catus - Ursus maritimus* | 11628 | 227 | 185 | 35 | 5.896E-25 | 328 | 41 | 5.936E-22 |
| *Felis catus - Zalophus californianus* | 5268 | 72 | 59 | 5 | 1.170E-03 | 89 | 5 | 7.070E-03 |
| *Mustela putorius - Neomonachus schauinslandi* | 12565 | 389 | 201 | 20 | 4.234E-06 | 665 | 38 | 1.918E-04 |
| *Mustela putorius - Odobenus rosmarus* | 12441 | 522 | 228 | 16 | 3.082E-02 | 694 | 54 | 7.614E-06 |
| *Mustela putorius - Panthera pardus* | 12334 | 323 | 222 | 21 | 3.696E-07 | 688 | 44 | 3.174E-08 |
| *Mustela putorius - Panthera tigris* | 11905 | 415 | 238 | 61 | 2.653E-36 | 780 | 94 | 1.266E-27 |

| | | | TreeShrink Outliers (small list) | | | TreeShrink outliers (large list) | | |
|---|---|---|---|---|---|---|---|---|
| **Species pair** | **Genes** | **Syntenic outliers** | **Total** | **Syntenic outliers** | **P-value** | **Total** | **Syntenic outliers** | **P-value** |
| *Mustela putorius - Puma concolor* | 10064 | 428 | 201 | 50 | 3.057E-25 | 677 | 75 | 6.000E-15 |
| *Mustela putorius - Suricata suricatta* | 10496 | 303 | 194 | 30 | 3.455E-14 | 906 | 56 | 2.985E-08 |
| *Mustela putorius - Ursus maritimus* | 12097 | 327 | 243 | 35 | 3.285E-16 | 776 | 58 | 8.841E-13 |
| *Mustela putorius - Zalophus californianus* | 5537 | 174 | 81 | 4 | 2.498E-01 | 300 | 11 | 3.424E-01 |
| *Neomonachus schauinslandi - Odobenus rosmarus* | 13241 | 568 | 171 | 21 | 1.364E-05 | 171 | 21 | 1.364E-05 |
| *Neomonachus schauinslandi - Panthera pardus* | 13160 | 357 | 188 | 22 | 7.913E-09 | 188 | 22 | 7.913E-09 |
| *Neomonachus schauinslandi - Panthera tigris* | 12635 | 413 | 215 | 67 | 9.955E-48 | 308 | 72 | 8.459E-42 |
| *Neomonachus schauinslandi - Puma concolor* | 10711 | 449 | 185 | 65 | 2.539E-43 | 288 | 74 | 1.210E-38 |
| *Neomonachus schauinslandi - Suricata suricatta* | 11126 | 328 | 190 | 25 | 3.098E-10 | 594 | 37 | 1.265E-05 |
| *Neomonachus schauinslandi - Ursus maritimus* | 12855 | 380 | 213 | 44 | 4.467E-25 | 303 | 48 | 5.249E-22 |
| *Neomonachus schauinslandi - Zalophus californianus* | 6052 | 163 | 61 | 9 | 3.119E-05 | 61 | 9 | 3.119E-05 |
| *Odobenus rosmarus - Panthera pardus* | 12988 | 538 | 209 | 28 | 3.912E-08 | 209 | 28 | 3.912E-08 |
| *Odobenus rosmarus - Panthera tigris* | 12447 | 613 | 243 | 74 | 5.050E-39 | 336 | 80 | 1.199E-33 |
| *Odobenus rosmarus - Puma concolor* | 10558 | 665 | 202 | 66 | 1.437E-30 | 309 | 82 | 1.373E-30 |
| *Odobenus rosmarus - Suricata suricatta* | 11012 | 521 | 210 | 29 | 1.881E-07 | 608 | 47 | 5.724E-04 |

| | | | TreeShrink Outliers (small list) | | | TreeShrink outliers (large list) | | |
|---|---|---|---|---|---|---|---|---|
| **Species pair** | **Genes** | **Syntenic outliers** | **Total** | **Syntenic outliers** | **P-value** | **Total** | **Syntenic outliers** | **P-value** |
| *Odobenus rosmarus - Ursus maritimus* | 12672 | 554 | 239 | 47 | 1.508E-18 | 327 | 52 | 3.018E-16 |
| *Odobenus rosmarus - Zalophus californianus* | 5973 | 351 | 68 | 7 | 1.027E-01 | 68 | 7 | 1.027E-01 |
| *Panthera pardus - Panthera tigris* | 12541 | 381 | 194 | 66 | 5.268E-52 | 265 | 69 | 1.357E-45 |
| *Panthera pardus - Puma concolor* | 10607 | 428 | 174 | 64 | 3.513E-45 | 262 | 72 | 6.301E-41 |
| *Panthera pardus - Suricata suricatta* | 10955 | 290 | 176 | 26 | 7.913E-13 | 582 | 34 | 1.129E-05 |
| *Panthera pardus - Ursus maritimus* | 12660 | 323 | 229 | 47 | 1.580E-29 | 319 | 50 | 1.141E-25 |
| *Panthera pardus - Zalophus californianus* | 5938 | 135 | 74 | 8 | 2.472E-04 | 74 | 8 | 2.472E-04 |
| *Panthera tigris - Puma concolor* | 10394 | 439 | 173 | 66 | 1.585E-46 | 316 | 80 | 3.867E-41 |
| *Panthera tigris - Suricata suricatta* | 10583 | 331 | 210 | 60 | 9.452E-42 | 676 | 74 | 2.804E-22 |
| *Panthera tigris - Ursus maritimus* | 12504 | 363 | 234 | 62 | 1.196E-42 | 408 | 69 | 5.657E-34 |
| *Panthera tigris - Zalophus californianus* | 5593 | 203 | 110 | 46 | 1.011E-38 | 159 | 50 | 5.493E-35 |
| *Puma concolor - Suricata suricatta* | 9020 | 359 | 178 | 60 | 1.957E-40 | 581 | 75 | 1.176E-20 |
| *Puma concolor - Ursus maritimus* | 10452 | 413 | 193 | 67 | 4.736E-46 | 355 | 78 | 1.725E-37 |
| *Puma concolor - Zalophus californianus* | 5164 | 209 | 96 | 37 | 8.113E-28 | 156 | 45 | 1.527E-27 |
| *Suricata suricatta - Ursus maritimus* | 10741 | 270 | 212 | 43 | 2.693E-27 | 695 | 57 | 6.877E-16 |

| | | | TreeShrink Outliers (small list) | | | TreeShrink outliers (large list) | | |
|---|---|---|---|---|---|---|---|---|
| **Species pair** | **Genes** | **Syntenic outliers** | **Total** | **Syntenic outliers** | **P-value** | **Total** | **Syntenic outliers** | **P-value** |
| *Suricata suricatta - Zalophus californianus* | 5123 | 132 | 94 | 15 | 1.088E-08 | 270 | 23 | 2.635E-07 |
| *Ursus maritimus - Zalophus californianus* | 5717 | 158 | 107 | 28 | 1.474E-20 | 159 | 29 | 1.239E-16 |
